# Sex differences in coronary artery disease and diabetes revealed by scRNA-Seq and CITE-Seq of human CD4+ T cells

**DOI:** 10.1101/2022.05.16.491900

**Authors:** Ryosuke Saigusa, Jenifer Vallejo, Rishab Gulati, Sujit Silas Armstrong Suthahar, Vasantika Suryawanshi, Ahmad Alimadadi, Jeff Markings, Christopher P. Durant, Antoine Freuchet, Payel Roy, Yanal Ghosheh, William Pandori, Tanyaporn Pattarabanjird, Fabrizio Drago, Coleen A. McNamara, Avishai Shemesh, Lewis L. Lanier, Catherine C. Hedrick, Klaus Ley

## Abstract

**Background:** Despite the decades-old knowledge that diabetes mellitus (DM) is a major risk factor for cardiovascular disease (CVD), the reasons for this association are only partially understood. Among the immune cells involved in CVD development, accumulating evidence supports the critical role of T cells as drivers and modifiers of this condition. CD4+ T cells are commonly found in atherosclerotic plaques. The activity and distribution of CD4+ T cell subsets differs between the sexes.

**Methods:** Peripheral blood mononuclear cells (PBMCs) of 61 men and women who underwent cardiac catheterization were interrogated by single cell RNA sequencing (scRNA-Seq, ∼200,000 cells) combined with 49 protein markers (CITE-Seq). Coronary artery disease (CAD) was quantified using Gensini scores, with scores above 30 considered CAD+ and below 6 considered CAD-. Four pairs of groups were matched for clinical and demographic parameters. To test how DM changed cell proportions and gene expression, we compared matched groups of diabetic and non-diabetic subjects. We analyzed 41,782 single CD4+ T cell transcriptomes for sex differences in 61 mostly statin-treated coronary artery disease patients with and without DM.

**Results:** We identified 16 clusters in CD4 T cells. The proportion of cells in CD4 cluster 8 (CD4T8, CCR2+ Em) was significantly decreased in CAD+, especially among DM+ participants. The proportions of cells in CD4T2, CD4T11, CD4T16 were increased and CD4T13 was decreased in CAD+ among DM+Statin+ participants. CD4T12 was increased in DM+ participants. In female participants, CD4T8, 12, and 13 were decreased compared to in male participants. In CD4 T cells, 31 genes showed significant and coordinated upregulation in both CAD and DM. The DM gene signature was partially additive to the CAD gene signature.

**Conclusions:** We conclude that CAD and DM are clearly reflected in PBMC transcriptomes and that significant differences exist between women and men and between subjects treated with statins or not.

## INTRODUCTION

Atherosclerosis is a chronic inflammatory disease of large and medium-sized arteries. People with type 2 diabetes mellitus (DM) have 2 to 4 times elevated risks of death and cardiovascular events compared to the general population.^1,2^ Despite the decades-old knowledge that DM is a major risk factor for cardiovascular disease (CVD), the reasons for this association are only partially understood. DM accelerates progression of atherosclerotic lesions and leads to defects in remodeling of plaques even after cholesterol reduction.^3^ Newer classes of drugs approved for glucose lowering in DM reduce CVD, but the mechanisms of this benefit do not appear to be explained by glucose lowering.^4^ Whereas the effects of glucose lowering on CVD outcomes are unclear, LDL-C lowering consistently reduces CVD risk associated with DM. Both type1 DM and type 2 DM are associated with greater vascular inflammation.^5^

Pre-menopausal women are significantly protected from CVD.^6^ Later in life, CVD risk in women catches up with that of men.^7^ Ultimately, myocardial infarctions and other manifestations of CVD become the #1 cause of death in women older than 85 years in the United States.^8^ The reasons for these sex differences in atherosclerosis and CVD^9^ are not well understood. Here, we focus on such differences, because CVD and its sequelae are a major public health problem.^10^

Among inflammatory and immune cell markers, the neutrophil–lymphocyte ratio is an easily obtained inflammatory biomarker that independently predicts cardiovascular risk and all-cause mortality^11^. Among the immune cells involved in CVD development, accumulating evidence supports the critical role of T cells as drivers and modifiers of this condition.^12^ CD4+ T cells are commonly found in atherosclerotic plaques. Some CD4 T cells recognize specific epitopes in apolipoprotein B (APOB)^13,14 (and Saigusa 2022 in press)^.

Early studies of DM focused on innate immune function. Recent studies suggest the adaptive immune system, especially T cells, also play a pivotal role in the pathogenesis of T2DM.^15^ A recent study demonstrated that activated CD4 T cells increased in the visceral adipose tissue of obese mice.^16^ These cells expressed PD-1 and CD153 and displayed characteristics of cellular senescence.^16^ It has also been shown that obesity induces MHC class II expression on adipocytes and thus activates CD4 T cells to initiate adipose tissue inflammation.^17^ These studies suggest CD4+ T cells may play an important role in obesity and obesity-induced insulin resistance.

Among adult humans, sex differences in lymphocyte subsets including T cells are described for multiple ethnic groups including Europeans, Asians, and Africans. Females have higher CD4+ T cell counts and higher CD4/CD8 ratios than age-matched males; whereas males have higher CD8+ T cell frequencies.^18^ The activity and distribution of CD4+ T cell subsets differs between the sexes. For example, naive CD4+ T cells from human females preferentially produce IFNγ upon stimulation, whereas naive T cells from males produce more IL-17.^19^

To elucidate the impact of CVD, DM and sex on CD4+ T cells, we investigated CD4+ T cells from high-quality frozen peripheral blood mononuclear cells (PBMCs) of 61 men and women with or without DM who underwent cardiac catheterization at the University of Virginia. These subjects are part of the Cardiovascular Assessment Virginia (CAVA) cohort.^20^ All cells were interrogated by targeted single cell RNA sequencing (scRNA-Seq) combined with 49 surface protein markers (CITE-Seq).

## METHODS

### Human Subjects

Subjects with suspected coronary artery disease (age range 40-80 years old) from the Coronary Assessment in Virginia cohort (CAVA) were recruited for study through the Cardiac Catheterization laboratory at the University of Virginia Health System, Charlottesville, Virginia, USA. All participants provided written informed consent before enrollment, and the study was approved by the Human Institutional Review Board (IRB No. 15328). Peripheral blood was obtained from these participants prior to catheterization. The study was approved by the Human Institutional Review Board (IRB No. 16017) at the University of Virginia.

### Quantitative Coronary Angiography (QCA)

Patients underwent standard cardiac catheterization with two orthogonal views of the right coronary artery and four of the left coronary artery according to accepted standards. QCA was performed using automatic edge detection from an end-diastolic frame. For each lesion, the frame was selected based on demonstration of the most severe stenosis with minimal foreshortening and branch overlap. The minimum lumen diameter, reference diameter, percent diameter stenosis, and stenosis length were calculated. Analysis was performed by blinded, experienced investigators. The Gensini score^21^ was used to assign a score of disease burden to each patient. Briefly, each artery segment is assigned a score of 0-32 based on the percent stenosis. The severity score for each segment was multiplied by 0.5-5, depending on the location of the stenosis. Scores for all segments were then added together to given a final score of angiographic disease burden. Score adjustment for collateral was not performed for this study. Subjects with Gensini score > 30 were classified as high CAD severity subjects and subjects with Gensini score < 6 were classified as low CAD severity subjects.

### Preparation of PBMC samples for CITE-seq

Blood from coronary artery disease subjects as well as subjects that had undergone cardiac catheterization to exclude CAD was drawn into BD K2 EDTA vacutainer tubes and processed at room temperature (RT) within one hour of collection. Whole blood in vacutainers were centrifuged at 400 x g for 10 minutes at RT to remove platelet rich plasma. PBMCs were isolated by Ficoll-Paque PLUS (GE Healthcare Biosciences AB) gradient centrifugation using SepMate-50 tubes (Stemcell Technologies Inc) following the manufacturer’s protocol. Trypan blue staining of PBMCs was performed to quantify live cell counts. PBMCs were cryopreserved in freezing solution (90% FBS/10% DMSO). PBMC vials were stored in Mr. Frosty (Thermo Fisher) for 48 hours at -80°C and were then stored in liquid nitrogen until used. To avoid batch effects, 8 samples each were processed on the same day, thawed in a 37°C water bath, centrifuged at 400 xg for 5 minutes and pellets resuspended in cold staining. All reagents, manufacturers and catalogue numbers are listed in **Table S1**. The viability and cell count of each tube was determined using the BD Rhapsody Scanner (**Table S2**). The tubes were then centrifuged at 400 xg for 5 minutes and resuspended in a cocktail of 51 AbSeq antibodies (2 μL each and 20 μL of SB, listed in **Table S3**) on ice for 30-60 minutes per manufacturer’s recommendations, washed and counted again. Of 65 samples investigated, 61 passed quality control (cell viability above 80%). Cells from each subjzect were sample tagged using Sample Multiplexing Kit (BD Biosciences) which contains oligonucleotide cell labeling, washed 3x, mixed, counted, stained with the 49 antibody mix, washed 3x again and loaded onto Rhapsody nanowell plates (4 samples per plate).

### Library preparation was completed according to BD’s recommendations

Pre-sequencing quality control (QC) was done using Agilent TapeStation high sensitivity D1000 screentape. Each tube was then cleaned with AMPure XP beads and washed in 80 % ethanol. The cDNA was eluted, a second Tapestation QC performed and diluted as needed. The samples were pooled and sequenced as recommended: AbSeq: 40,000 reads per cell, mRNA: 20,000 reads per cell, sample tags: 600 reads per cell on Illumina NovaSeq using S1 and S2 100 cycle kits (Illumina) (67×8×50 bp). FASTA and FASTQ files were uploaded to the Seven Bridges Genomics pipeline, where the data was filter to generate matrices and csv files. This analysis generated draft transcriptomes and surface phenotypes of 213,515 cells (496 genes, 51 antibodies). After removing multiplets based on sample tags and undetermined cells, 175,628 cells remained. Doublet Finder (https://github.com/chris-mcginnis-ucsf/DoubletFinder) was used to remove additional doublets, leaving 162,454 cells. 961 CD4+ T cells were removed because they looked like biological doublets with myeloid cells.^22^ CD4+ T cells were defined as CD19- CD14- CD16- CD3+ CD4+ CD8-. 40,821 CD4+ T cells were detected. All antibody data were CLR (centered log-ratio) normalized and converted to log2 scale. All transcripts were normalized by total UMIs in each cell and scaled to 1000.

### Thresholding

Each antibody threshold (**Table S4**) was obtained by determining its signal in a known negative cell or by deconvolution of overlapping normal distributions (we used the function ‘normalmixEM’ to deconvolute the overlapping distributions in the package ‘mixtools’). To identify the thresholds, ridgeline plots for each antibody in each main cell type were used to set the best threshold (**Figure S1**).

### Clustering

We use UMAP (Uniform Manifold Approximation and Projection) dimensionality reduction in order to project the cells onto the 2D space. We selected UMAP as our choice of algorithm because it focuses on capturing the local similarities while at the same time preserving the global structure of the data. We ran the UMAP algorithm over the top 20 principal components given by Harmony. In order to cluster the data, we use the standard Louvain clustering algorithm with the parameters resolution set to 0.15 and random seed set to 42 in order to ensure the reproducibility of the results. Before running the Louvain clustering, we filtered out antibodies that are not expressed (CD19 and CD8 for CD4+ T cells).

### Comparing Gene Expression among Participant Types

To determine differential expression (DE), we used the Seurat package in R with no thresholds over avg_logFC, minimum fraction of cells required in the two populations being compared, minimum number of cells and minimum number of cells expressing a feature in either group and filtered for adjusted p<0.05

### Comparing Cell Proportions

To find changes in proportions, we identified the cell numbers for each participant in each cluster, calculated the log-odds ratio p/(1-p), where p is the proportion of cells, followed by pairwise comparison.

### Random Forest Model

A machine learning (ML) approach was implemented to identify the genes with the highest capability to distinguish between disease groups. To accomplish this goal, the Random Forest (RF) model^23,24^ was trained with the normalized gene expression from 1000 randomly selected cells from each condition and variable importance scores of the genes were calculated. This procedure was repeated for 15 iterations and importance scores in each iteration were scaled to the 0-100 range for a better comparisons. A higher score is considered as a higher power in classifying the groups.

## RESULTS

### Population and cells

The 61 CAVA participants initially selected for the current study were aged 42-78 years, mostly non-hispanic Whites. Matched quartets were selected based on sex, Gensini score, DM status, and statin treatment: 17 Diabetic men with or without CAD, 17 Non-diabetic men with or without CAD, 16 Women with or without CAD. All these patients were statin-treated. In addition, we studied a group of 10 matched non-diabetic men with or without CAD, not statin-treated (statistical summary of clinical parameters is in **Table S5A-E**). Women and diabetic men without statin treatment were not found in this cohort. Categorical variables were compared by Chi-square test and continuous variables by Mann-Whitney test. PBMC tubes were shipped from the central repository on liquid N_2_, thawed and processed according to standard operating procedures, resulting in 91±4% cell viability (**Table S2**). To avoid batch effects, all cells were hash-tagged for multiplexing, with 4 samples run per 250,000-well plate (total of 16 plates). The pooled cells were labeled with 49 titrated oligonucleotide-tagged mAbs (**Table S3**). After quality controls and three-stage doublet removal, 140,610 single cell transcriptomes from 61 WIHS participants were successfully analyzed. Among them, 41,782 CD4 T cells were identified.

### Surface marker expression

In combined protein and transcript panel single cell sequencing, non-specific binding contributes to the antibody signal, in part, because Fc block is not complete. Additional background is the consequence of unbound oligonucleotide-tagged antibody remaining in the nanowell that will be amplified and sequenced. To account for all sources of background, we established a threshold for each antibody, using ridgeline plots separately for CD4+ T cells, CD8+ T cells, B cells and monocytes. This yielded thresholds for 34 markers used (**Table S4**). 15 were not thresholded, because they showed continuous expressions without clear threshold.

### CD4 T cell identification

CD4 T cells were identified as CD19-CD3+CD4+CD8-cells. CD4 T cells underwent Louvain clustering, and gates were overlaid and used in UMAP figures (**Figure 1A**). Initially 17 clusters were identified in CD4 T cells, but cluster 9 was identified as likely doublets between CD4 T cells and myeloid cells. Among 16 CD4 T cell clusters, we identified 3 naïve clusters, 1 Th1, 1 Th2, 1 Tfh and 1 Treg cluster, based on the heatmap of the informative surface marker (**Figure 1B**). Other clusters were identified as central memory (Cm) and 4 clusters as effector memory (Em): PDL1+, PD1+, CCR2+ and NK-like. One cluster of Em cells expressed CD45RA (Emra). Cluster 12 was MMP9+ and cluster 16 was PDCD1+. Violin plots for the expression of all surface markers in each cluster are shown in **Figure S2**.

**Figure 1.**
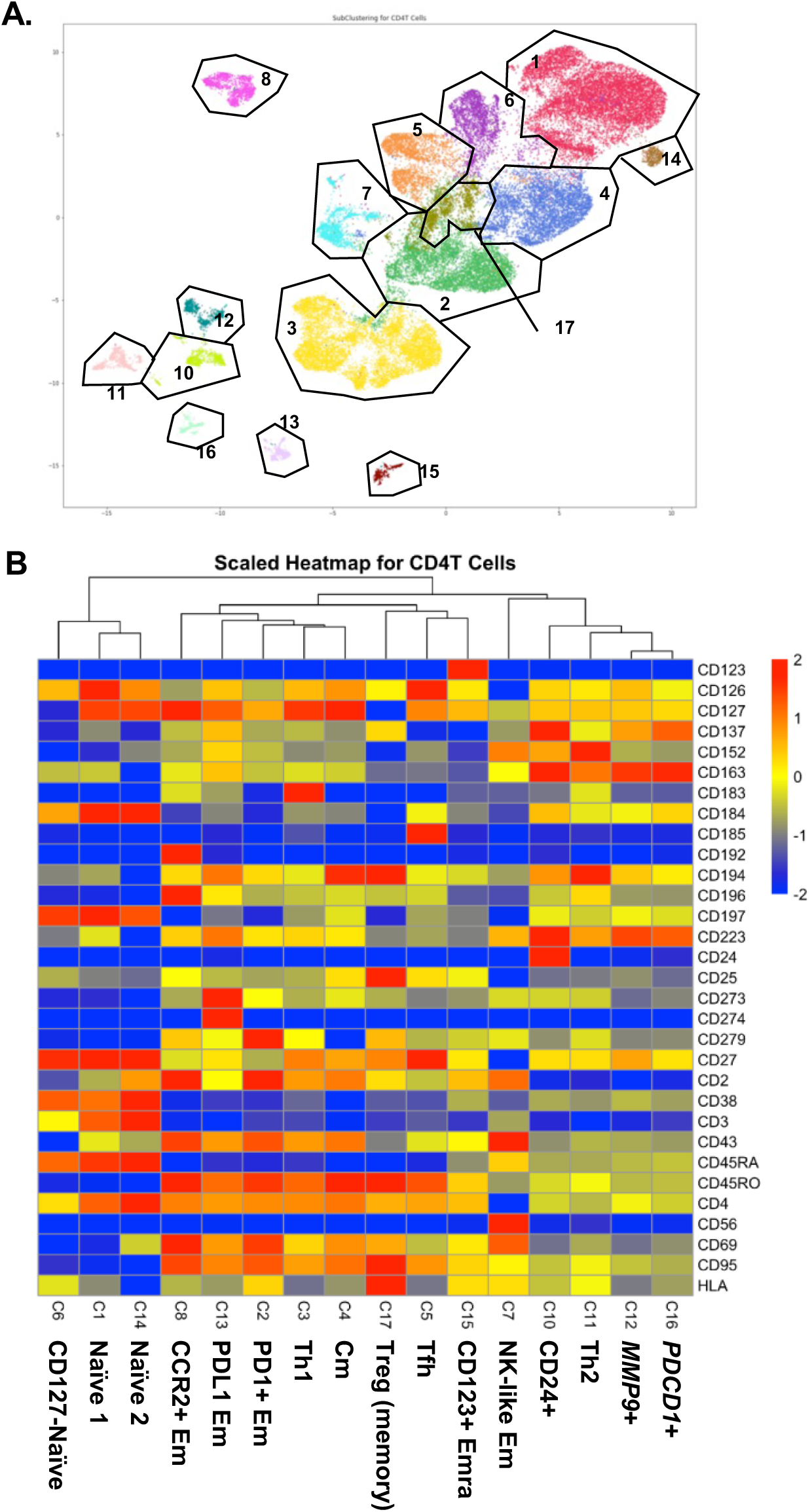
**A**, CD4 T cells were UMAP-Louvain-clustered by all non-negative surface markers. CD4 T cells formed 16 clusters, cluster numbers indicated. **B**, Scaled heatmaps of antibody expression in each subcluster, cluster numbers corresponding to panel A, cluster names indicated.

### Transcriptomes

Next, we analyzed the transcriptomes of each single cell (**Supplemental Excel File 1**). We tested gene expression of each cluster against all other clusters in CD4+ T cells (**Figure 2A** and **Supplemental Excel File 2**), using Seurat to report the data as log2 fold-change (logFC). The transcriptomes (**Figure 2**) confirmed the identity of the CD4+ T cell clusters identified by CITE-Seq (**Figure 1**) and expanded phenotype information.

**Figure 2.**
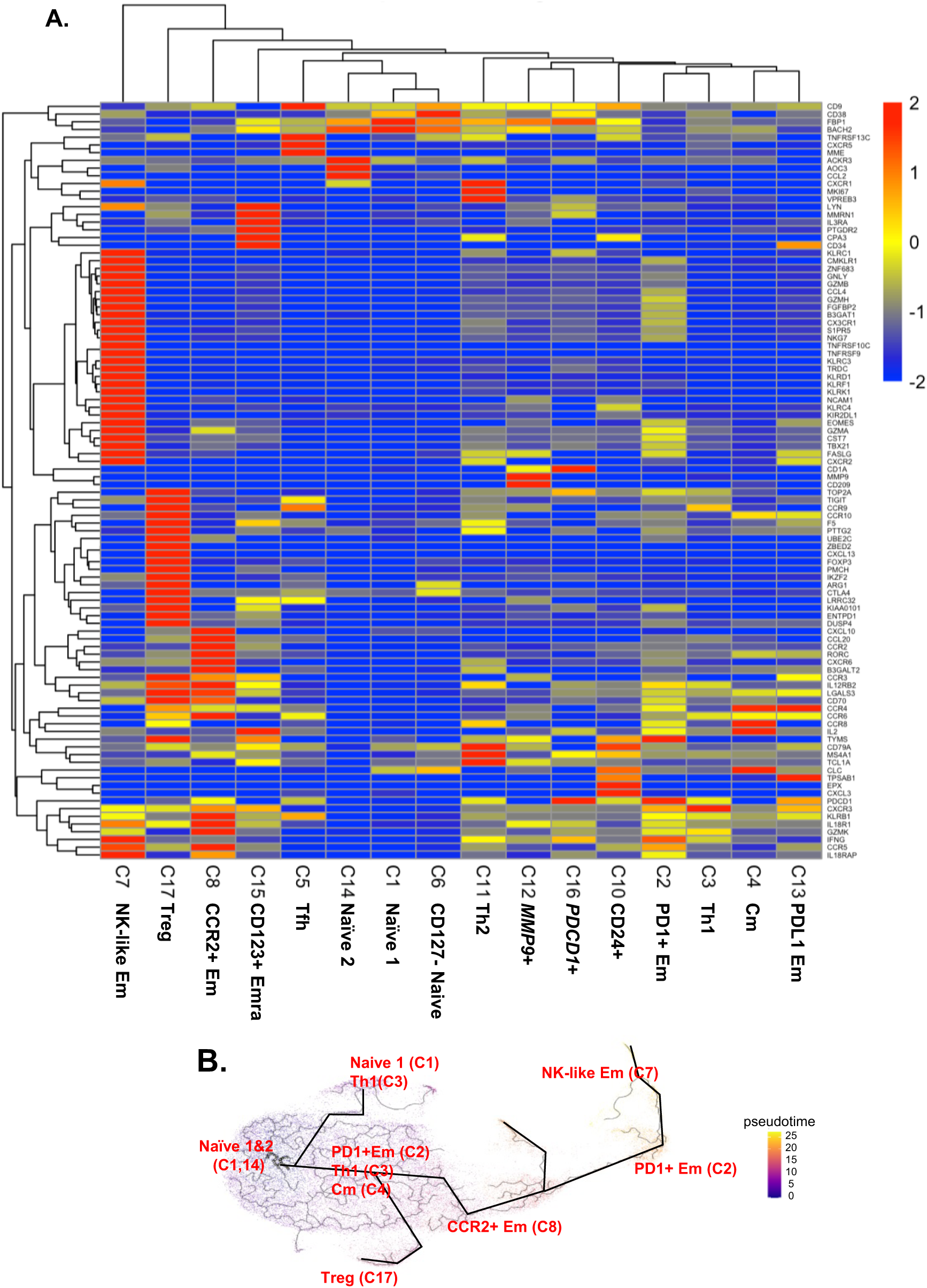
**A**, Scaled heatmaps of differentially expressed genes (filtered on the basis of adjusted p-val < 0.05, avg_log2FC > 0, and pct.1(percentages of cells expressing each gene in each cluster / those of cells expressing each gene in all the other clusters > 2.5) in each subcluster. **B**, Trajectory analysis by monocle 3. The starting node was defined as the naïve clusters 1 and 14. Pseudotime indicated by color (blue to yellow).

Some CD4 T cells are cytotoxic.^12^ Here, we found significant overexpression of cytotoxic genes like *GZMB, GZMH* and *NKG7*, also *FGFBP2*, and *GNLY* in the NK-like Em cluster 7. These genes were also higher in some cells in cluster 2. *KLRF1, KLRK1, KLRD1, CX3CR1, ZNF683* were unique to cluster 7. Some Treg signature genes including FoxP3, *LGLS3, TIGIT*, and *CCR3* were upregulated in the Treg cluster 17. CD4 T cells in cluster 8 (CCR2+) also expressed higher *CCR2* and *CCR6* transcripts, and also *KLRB1, GZMK*, and *RORC* gene, suggesting that this cluster contains Th17 cells. 2 Naïve clusters (cluster 1 and 14) expressed similar gene patterns, except higher *ACKR3, AOC3* and *CCL2* in cluster 14 but not cluster 1. Tfh (cluster 5) expressed the *CXCR5* gene, the classical chemokine receptor characteristic for Tfh cells.^25^ Clusters 1, 6 and 14 expressed low levels of *KLRB1*.

To test how these CD4 T cell subsets might be related, we conducted a trajectory analysis by Monocle3 (**Figure 2B**). This showed that the naïve CD4+ T cells in cluster 1 and 14 split into Th1 cells (cluster 3) and Treg cells (cluster 17). From a downstream node containing cluster 2 (PD1+ Em), central memory (Cm, cluster 4) and some cluster 3 Th1 cells, the CCR2+ Em (cluster 8) emerged. Finally, PD1+ Em (cluster 2) gave rise to NK-like Em cells (cluster 7).

### CD4+ T cell subset abundance

We asked whether, and if so, how, DM, CVD and sex affected the abundance (cell number) of CD4+ T cell subsets expressed as log odds ratios (Cell counts are in **Supplemental Excel File 3**). We compared the log-odds ratios for each of the CD4T cell clusters (**Figure 3**). The proportion of CCR2+ effector memory CD4 T cells (cluster 8) was significantly (p<0.01) lower in subjects with CAD than in those without CAD. This was mainly driven by a strong decrease in diabetic men with statin treatment (**Figure 3A**). CCR2 has been reported to be involved in Treg cell recruitment,^26^ but this role has not been shown in atherosclerosis. Women showed a significantly lower proportion of cluster 8 than men (**Figure 3A**).

**Figure 3.**
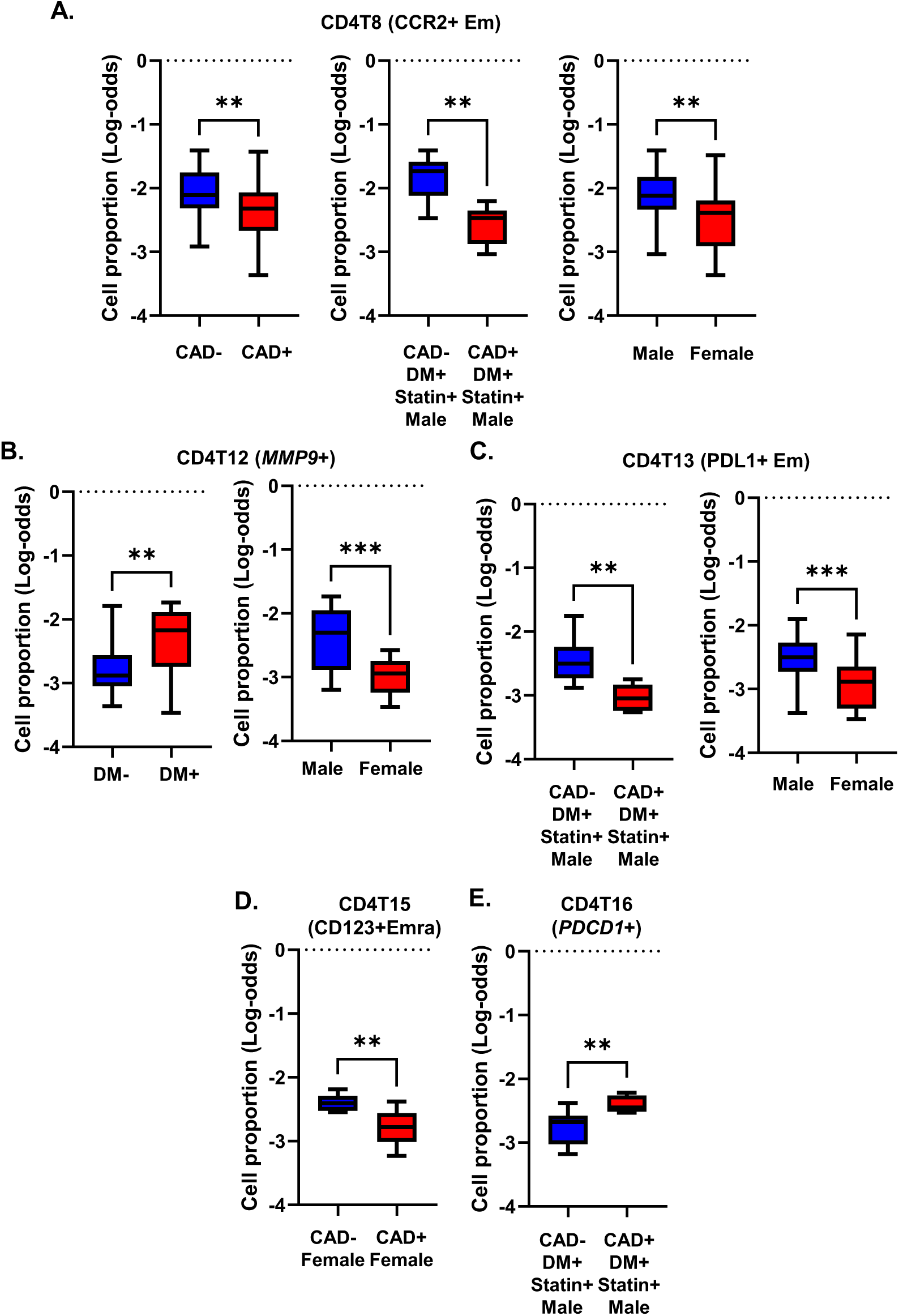
Cell proportions in men and women with or without CAD and/or DM. **A-E**, Proportions of cells in each cluster calculated as percentage of the parent cell type as indicated in the title of each panel. Clusters with significant differences (**, p<0.01) in cell proportions (by log odds ratio) are shown with means and standard error of the mean (SEM).

The proportion of *MMP9*+ CD4 T cells (cluster 12) was significantly higher in subjects with DM, regardless of CVD status (**Figure 3B**). This cluster was significantly less abundant in women than men. The proportion of PDL1+ effector memory CD4 T cells (cluster 13) was significantly elevated in males compared to females. This was mainly driven by a high abundance of these cells in males with DM and without CVD (**Figure 3C**). CD4T15 (CD123+Emra) was significantly decreased in women with CVD (**Figure 3D**). CD4T16 (*PDCD1*+) was increased in CAD+ in diabetic men with statin treatment (**Figure 3E**). Interestingly, cluster14 (Naïve 2) was observed only in non-diabetic men, regardless of CAD status (**Figure 4**).

**Figure 4.**
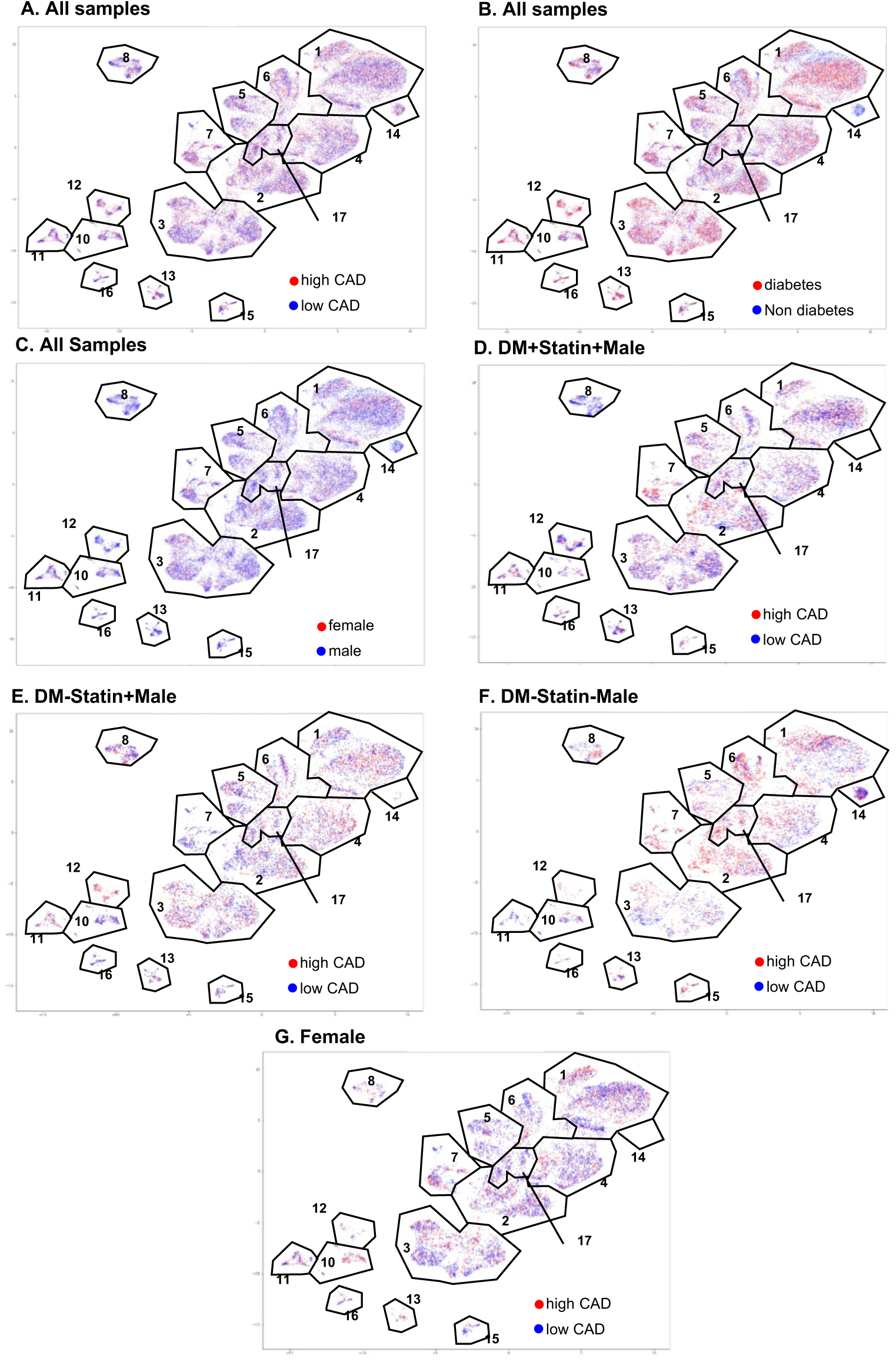
CAD, DM and sex features projected onto CD4+ T cell UMAPs. **A**, high CAD (red), and low CAD (blue) projected to UMAPs from all the samples. **B**, DM (red), and nonDM (blue) projected to UMAP from all the samples. **C**, female (red) and male (blue) projected to UMAP from all the samples. **D-G**, high CAD (red) and low CAD (blue) projected to UMAP from DM+statin+male samples (**D**), DM-statin+male samples (**E**), DM-statin-male (**F**), and all the females (**G**).

### Differentially expressed genes by disease status

In all CD4T cells, several genes including *TCF7, LTB*, and *GNAI2* were upregulated in CAD+ subjects (**Figure 5A**). In DM, *TCF7* and *GNAI2* were also upregulated, in addition to *LAIR2* and *IFITM3*. The known atherosclerosis gene *IL32*^27^ was downregulated in DM+ compared to non-DM participants (**Figure 5B**). More systematic investigation of the overlap showed that 31 genes were significantly upregulated in both CAD and DM (**Table S6** for the genes expressed in at least 20% of the higher expressing cell type (pct.1 > 0.2).

**Figure 5.**
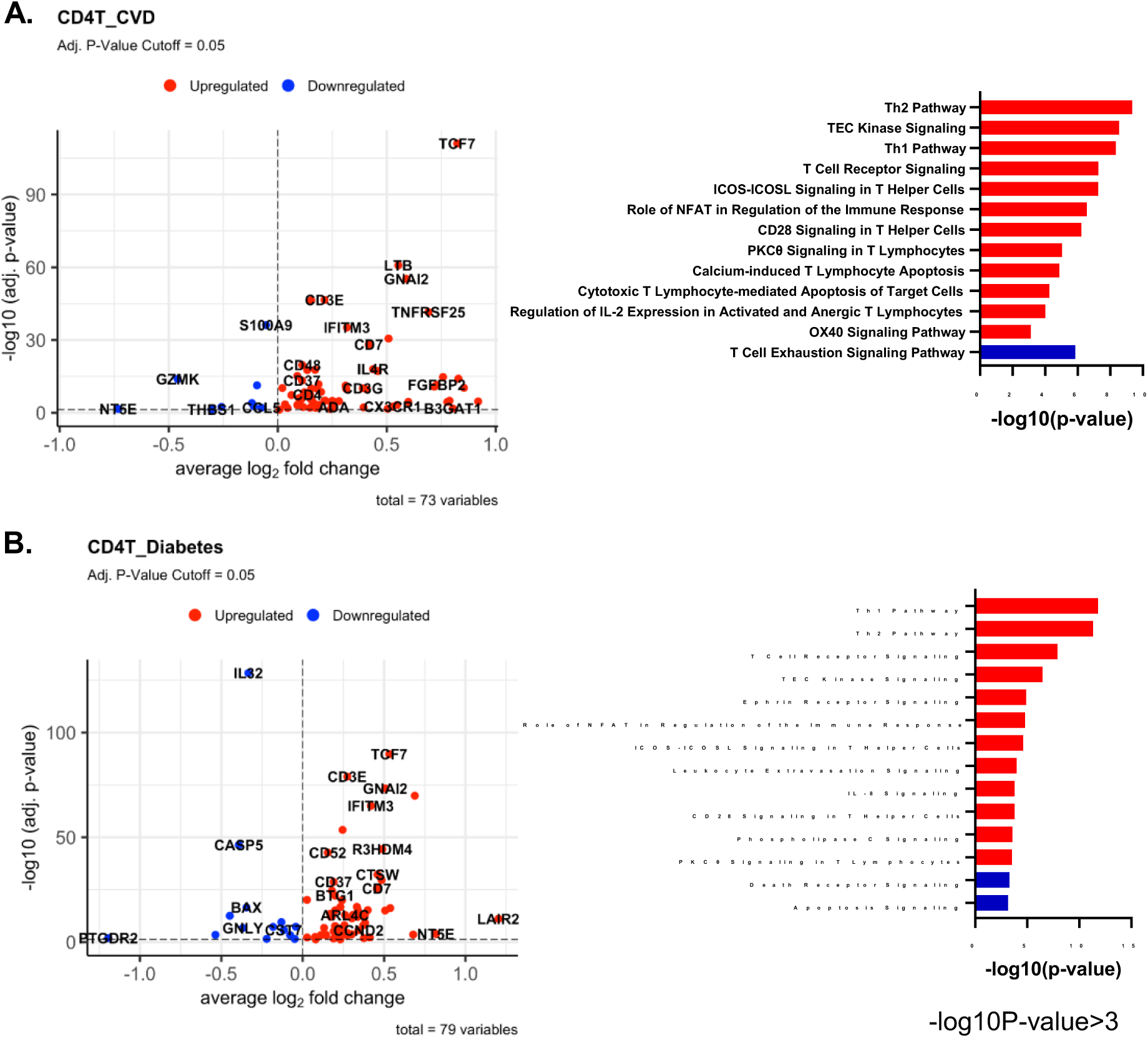
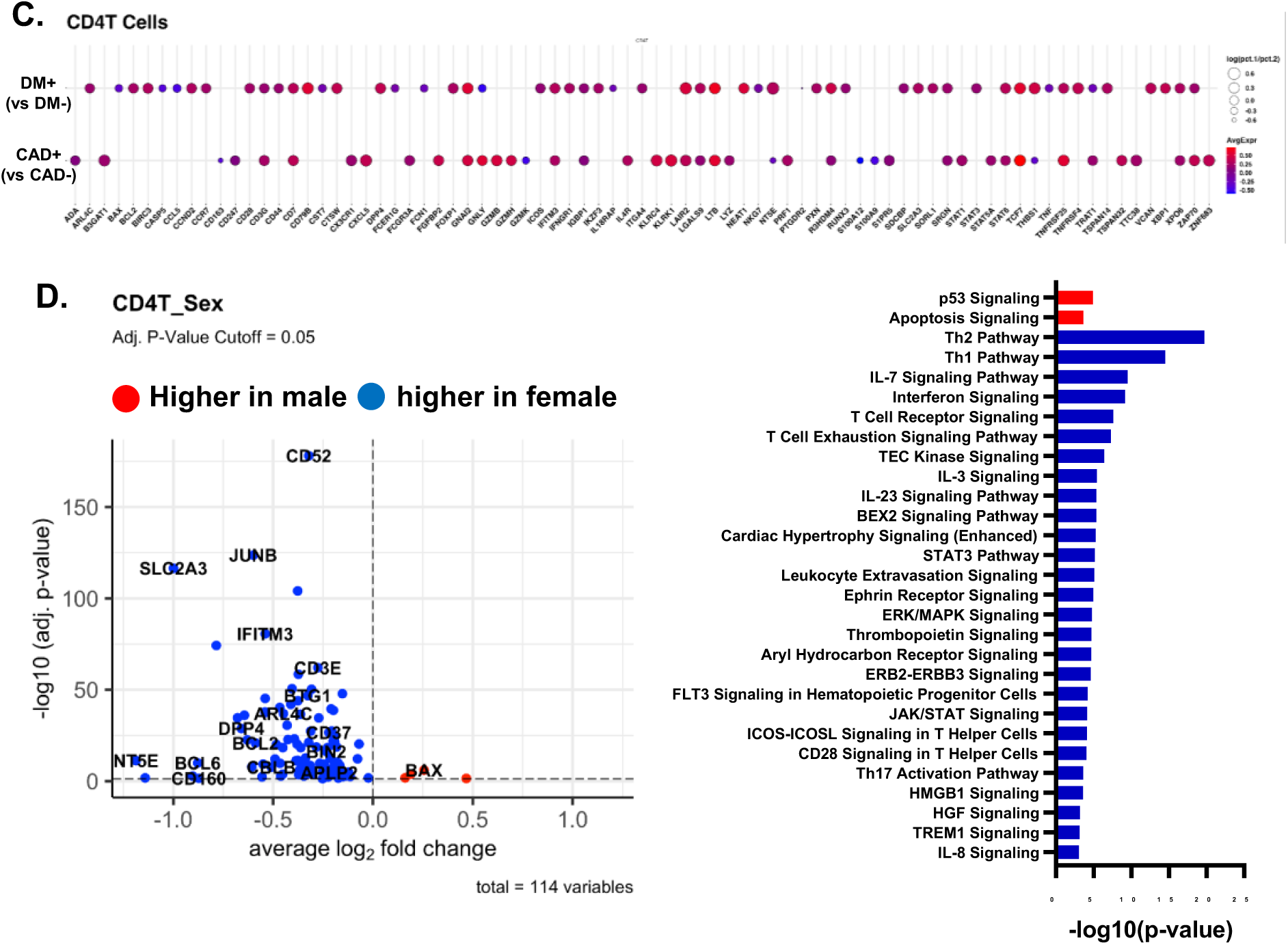
**A, B**, Volcano plots comparing gene expression in single cells and Ingenuity Pathway analysis (-log10 p-value>3) between CAD high vs CAD low (**A**) and between Diabetes+ vs DM- (**B**) in all CD4 T cells. Red bar means positive Z-score, and Blue bar means negative Z-score. **C**, Dotplots of differentially expressed genes between CAD+ vs CAD-, and between DM+ vs DM- in all CD4 T cells. The thresholds set for the plots were adjusted p-value <0.05, avg.Log2FC>0 or <0, and pct.1 > 0.2. The size of dots represents log(pct.1/pct.2), where pct.1 is the proportion of cells expressing each gene in DM or CAD+ and pct.2 is the proportion of cells expressing each gene in DM- or CAD-. **D**, Volcano plots comparing gene expression in single cells and Ingenuity Pathway analysis (-log10 p-value>3) between male and female in all CD4 T cells. Red bar means positive Z-score, blue bar means negative Z-score.

TCF7 has been reported to have a role in DM,^28,29^ but the function of TCF7 has not been reported either in CAD and or in DM. G-protein alpha-i2 coded by GNAI2 is related to CXCR5 signaling, which is thought to induce Hippo/YAP-dependent DM-accelerated atherosclerosis.^30^ This G-protein is important for T lymphocyte chemokine receptor signaling. Local chemoattractants regulate the movement of CD4 T cells.^31^ Specifically, GNAI2 was found to be required for chemokine-mediated arrest of rolling leukocytes.^32,33^ Volcano plots of differentially expressed genes (DEGs) for each cluster and dotplots of the common DEGs in CAD and DM are shown in **Figure S3 and S4**, respectively; underlying data in **supplemental Excel File 4 and 5**.

### Pathway analysis

In CD4 T cells, several pathways like “Th1”, “Role of NFAT in the Immune Response and T cell receptor signaling” were upregulated in both CAD+ and DM+ subjects (**Figure 5A, B and C**). Nuclear factor of activated T cells (NFAT) is a transcription factor with a multidirectional regulatory function. NFAT-centered signaling pathways play important regulatory roles in the progression of atherosclerosis. NFAT is involved in vascular smooth muscle cell phenotypic transition and migration, endothelial cell injury, macrophage-derived foam cell formation, and plaque calcification.^34^ Only in CAD, but not in DM, the “Cytotoxic T Lymphocyte-mediated Apoptosis of Target cells” pathway was highly upregulated, and “T cell exhaustion signaling” pathway was downregulated. In DM, “IL-8 signaling”, “Integrin signaling” and “Ephrin receptor signaling” pathways were highly and uniquely upregulated. Circulating IL-8 levels are increased in patients with type 2 DM. IL-8 is associated with worse inflammatory and cardio-metabolic profiles.^35^

We also applied pathway analysis to detect systematic sex differences (**Figure 5D**). Many genes including *CD52, JUNB, IFITM3* and *SLC2A3* were more highly expressed in females than males. This resulted in significant enrichment for the Th2, Th1, IL-7 signaling and interferon signaling pathways in females. Only 2 pathways, p53 signaling and apoptosis signaling, were enriched in men.

### Random Forest Analysis

To identify the genes with the highest capability to distinguish between disease groups, we used the random forest machine learning (ML) approach. To decrease the covariants, we focus on the participants treated with statins, the majority of our subjects. *TCF7* was the most important gene to separate CAD from non-CAD participants (**Figure 6A**), regardless of DM status (**Figure 6B, C**). The bottom panel of **Figure 5A-C** shows overlaid ridge plots for *TCF7, IL32, LTB, CD52*, and GNAI2. In DM, *TCF7, S100A9, IL32* and *KLF2* were overexpressed in CD4+ T cells from subjects with CAD (**Figure 6B**). In non-DM subjects, *TCF7, CD52* and *GNAI2* were overexpressed in CAD (**Figure 6C**). We correlated the gene ranks in DM and nonDM (**Figure 6D**), which showed that *S100A9* was more important in DM, and *IFITM3, FOSB*, and *CCL5* were more important in nonDM.

**Figure 6.**
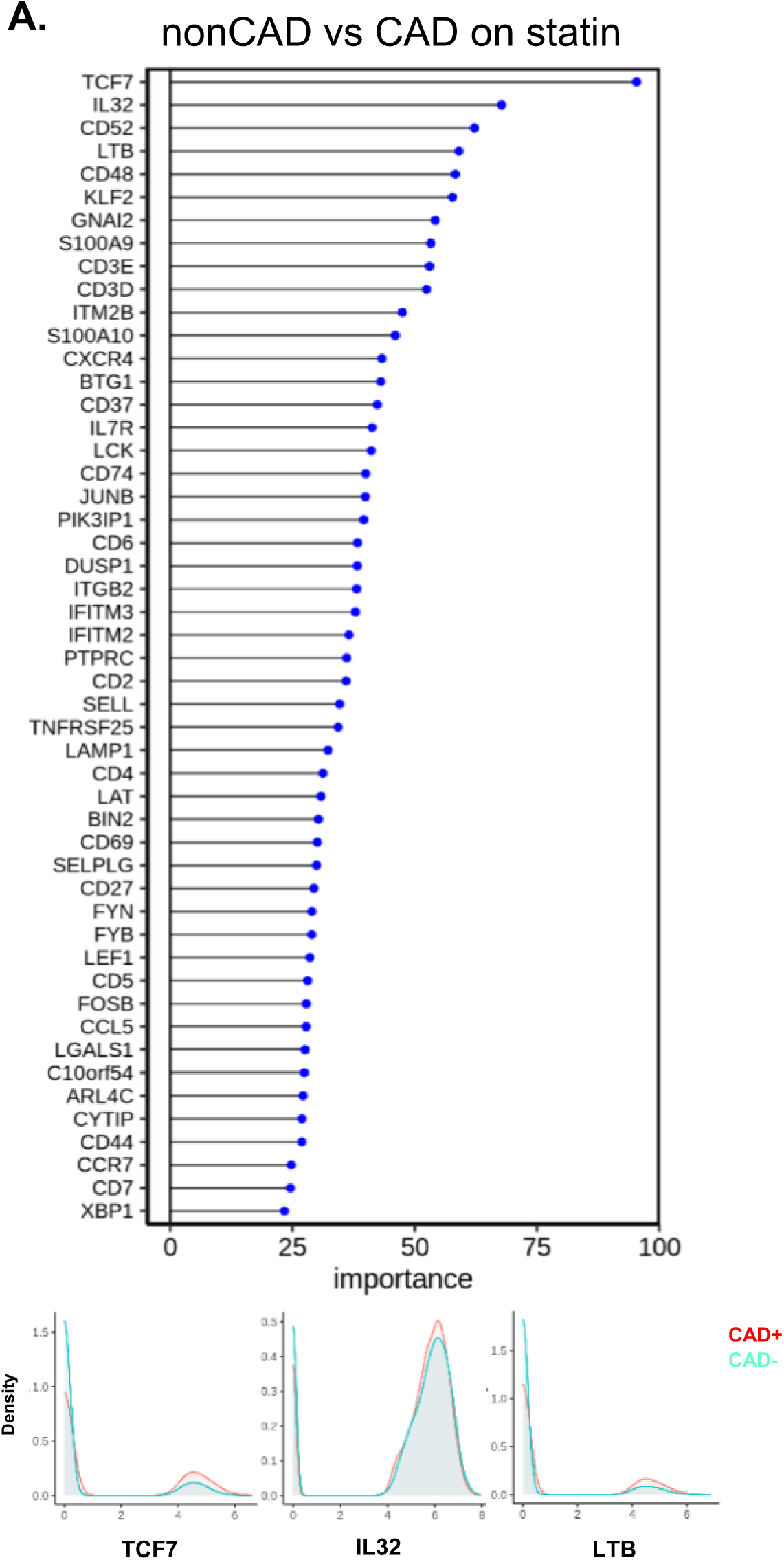
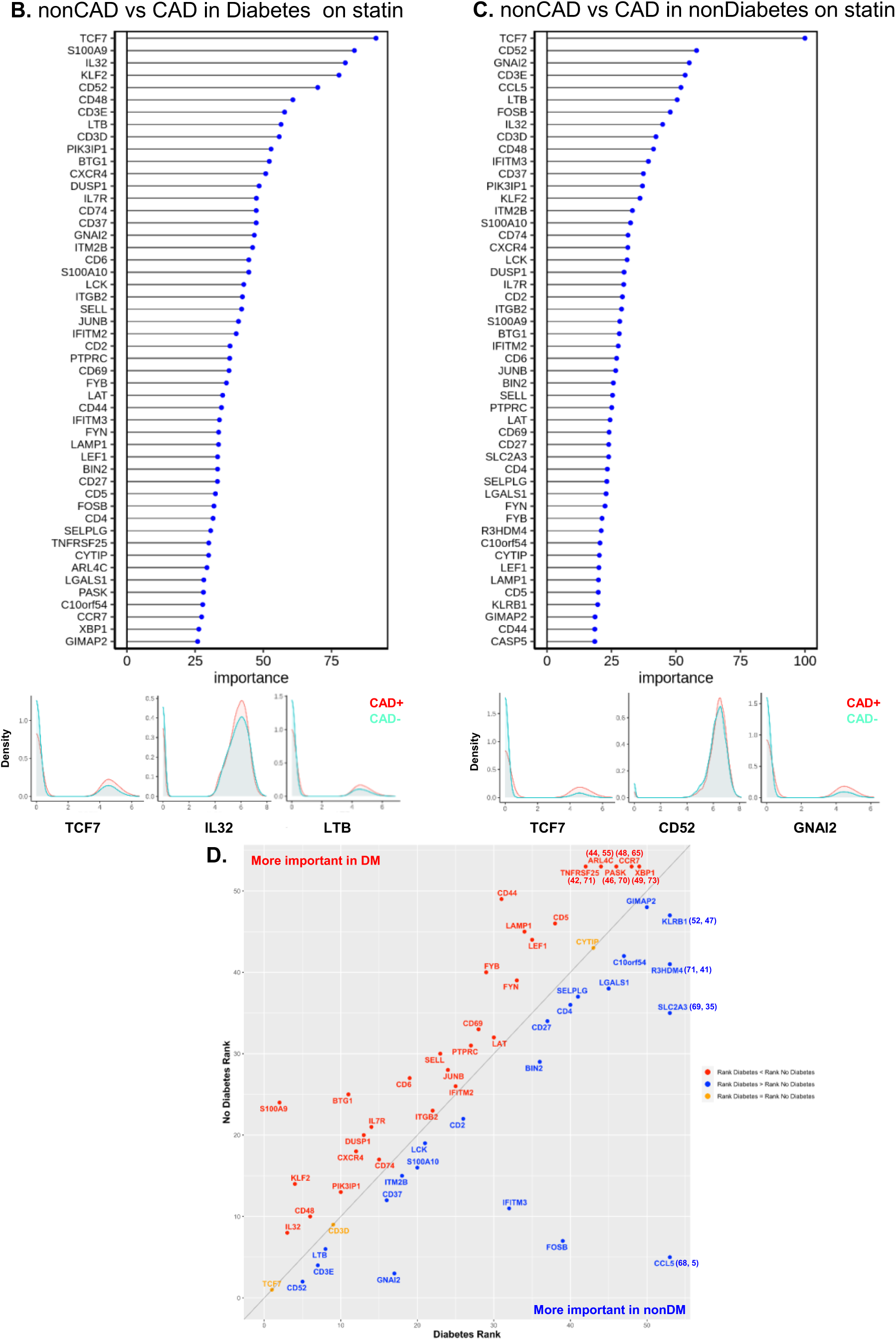
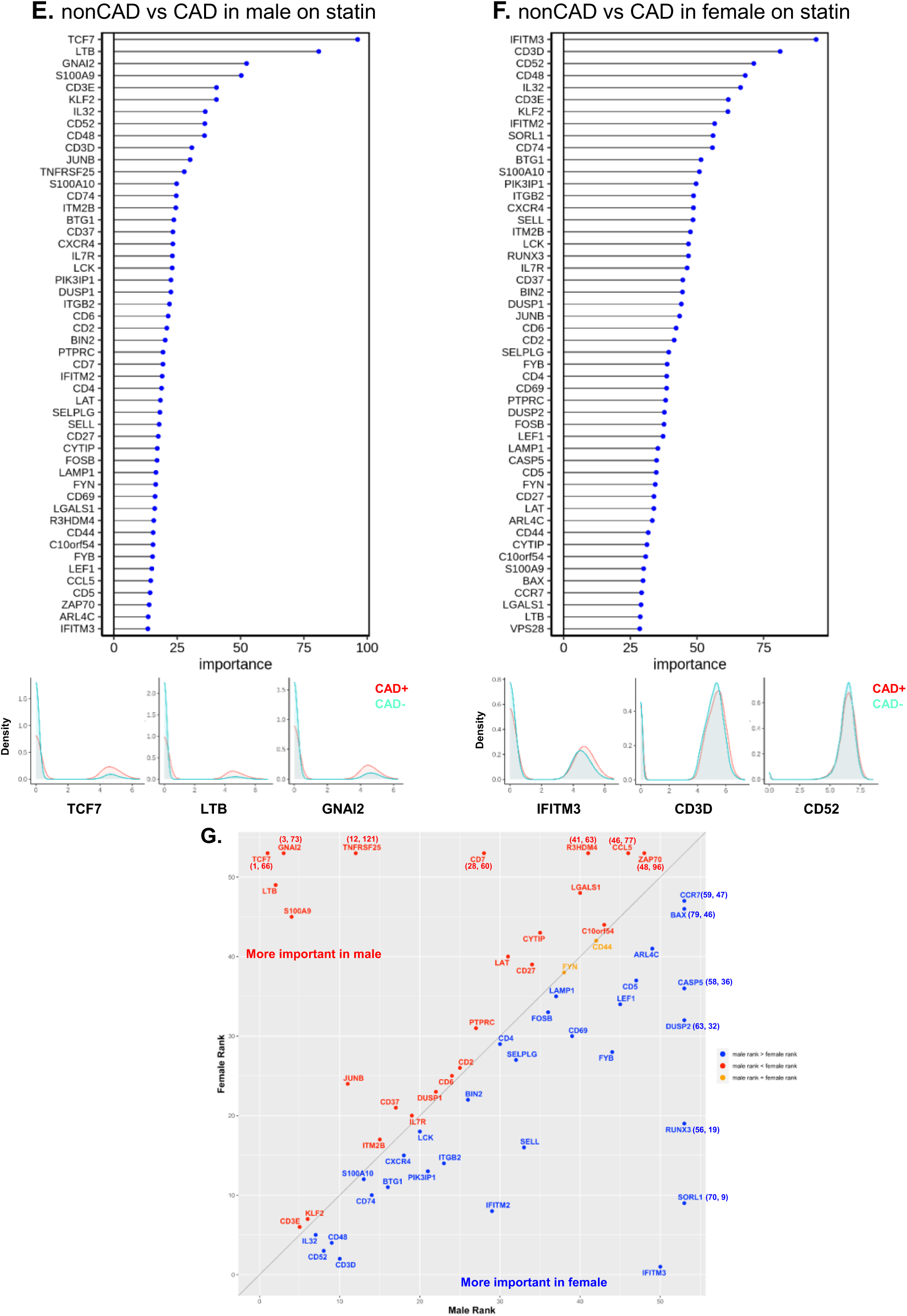
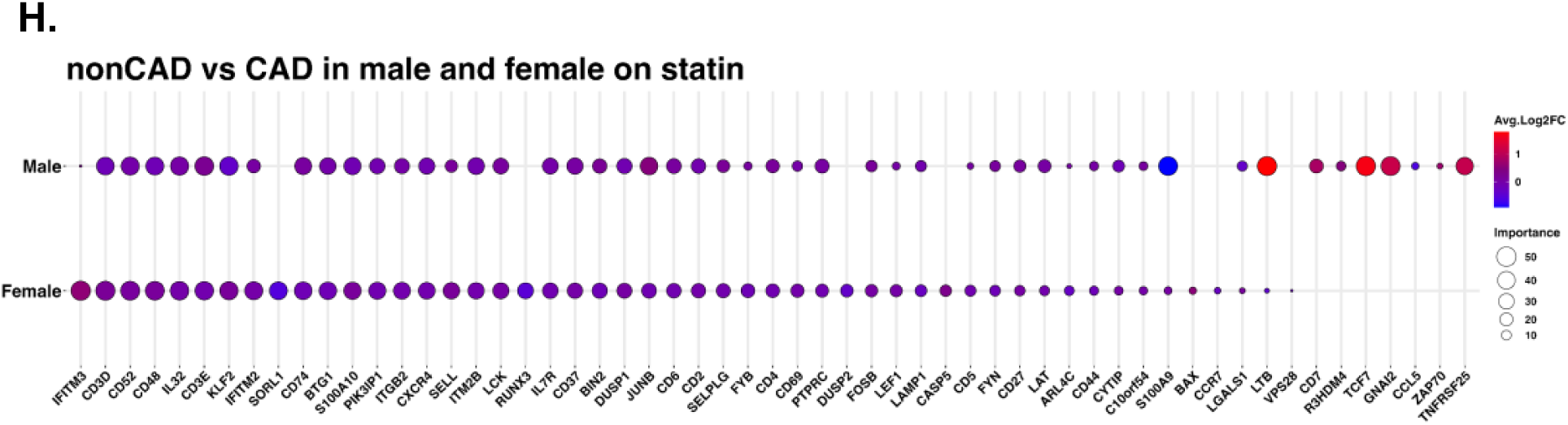
Random forest analysis. **A**, Line with dot plots which show the feature importance of each gene in the comparison between nonCAD vs CAD. Below, representative density plots are shown. **B, C, E, F**, line with dot plots for the feature importance of each gene in the comparison between nonCAD vs CAD in DM (**B**), in non DM (**C**), in males (**E**), and in females (**F**). At the bottom of each plot, representative density plots are shown. **D, G**, Correlation of the ranks of the important genes for CAD in DM (x-axis) vs nonDM (y-axis), and males (x-axis) and females (y-axis). Red dot, rank in DM < rank in nonDM; blue, rank in DM> rank in nonDM; yellow, rank in DM = rank in nonDM in **D**. Red dot, rank in male < rank in female; blue, rank in male > rank in female; yellow, rank in male = rank in female in **G**. When a gene ranked outside 50, an actual rank was shown. **H**, Dotplot which showed the importance of genes for CAD in male and female side by side. Average log2 fold change (log2FC) for male to female comparison.

Next, we investigated the driver genes for CAD in males and females separately (**Figure 6E, F**). Interestingly, *TCF7* was important in male, but not in female participants with CAD. *IFITM3* was the most important gene to separate CAD from low-CAD in women, but was ranked in 50^th^ in male. This gene encodes interferon-induced transmembrane protein 3. The discrepancies in gene expression in CD4+ T cells between women and men with and without CAD are striking and unexpected. CD4+ T cells from CAD+ men significantly overexpress *TCF7, LTB* and *GNAI2* (**Figure 6E**, lower panels), whereas the top genes in women are *IFITM3, CD3D* and *CD52* (**Figure 6F**, lower panels). We correlated the gene ranks in male and female (**Figure 6G**), which showed that *TCF7, LTB*, and *GNAI2* was more important in male, and *IFITM2, IFITM3*, and *SORL1* were more important in female. For those genes ranked in the top 50 in both men and women, we also constructed a side-by-side comparison of the relative importance (**Figure 6H, G**).

### Interaction Analysis

CD4 T cells interact with myeloid cells in the context of antigen presentation^36^, co-stimulation, co-inhibition and for inflammatory receptor-ligand interactions for cytokines and chemokines. We identified classical, nonclassical and intermediate monocytes as well as dendritic cells. Such interactions are known to be important for antigen presentation. T cells can also form biological doublets with myeloid cells in vivo.^22^ Here, we focus on predicted interactions with p<0.01, with the communication probability coded by color from low (blue) to high (red). All CD4+ T cell clusters from subjects with CVD showed interactions between LGALS9 and CD44 (**Figure 7A**), and all except cluster 14 showed this interaction in DM+ subjects (**Figure 7B**). Galectin-9 (coded by *LGALS9* gene) is a crucial regulator of T-cell immunity by inducing apoptosis in specific T-cell subpopulations associated with autoimmunity and inflammatory disease.^37^ About half of the CD4+ T cell clusters showed L-selectin (SELL) interaction with PSGL-1 (SELPLG) in both CVD+ and DM+ subjects (**Figure 7A, B**). Multiple studies have provided evidence supporting a key role for SELP and SELPLG in atherosclerotic lesion formation, thrombosis, and arterial wall changes.^38,39^ The SELL-SELPLG interaction was stronger in males than in females (**Figure 7E, F**). There was no male-female difference in LGALS9-CD44 interaction between males and females. Uniquely, only Tregs (cluster 17) showed CTLA4 interaction with CD86 in both males and females. In non-CVD and non-DM subjects (**Figure 7C, D**), the interaction pattern of CD4+ T cells with myeloid cells was similar, but significant interactions were fewer. The CTLA4 interaction with CD86 in Tregs was preserved.

**Figure 7.**
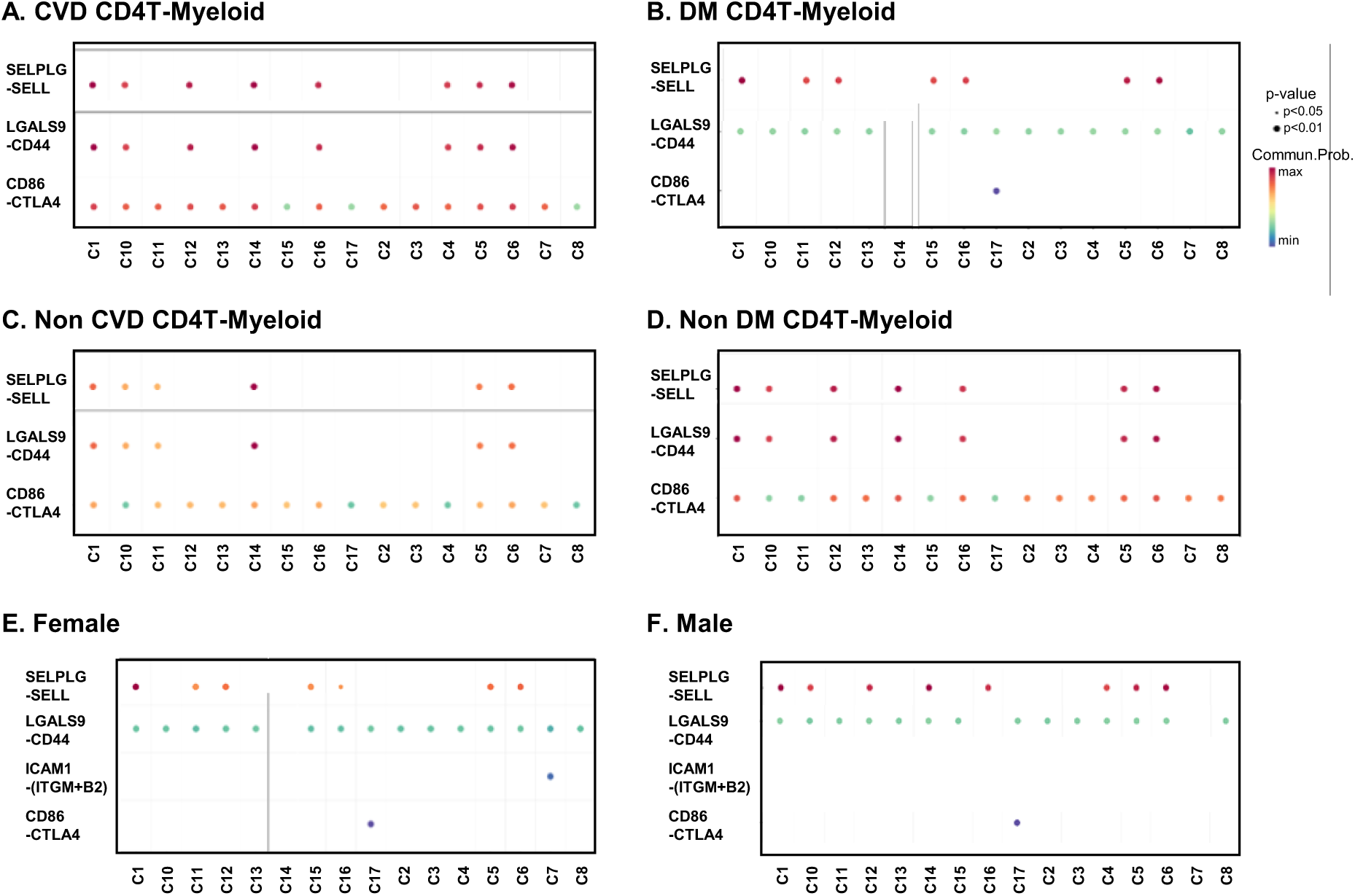
Bubble plots of interaction analysis between each CD4T cell cluster and myeloid cells. The color bar indicates communication probability. The size of bubbles indicates p-value.

## DISCUSSION

It is well known that coronary artery disease and its complications including myocardial infarction manifest differently in men and women.^40^ Plaque failure in women is more commonly attributed to erosion, whereas rupture is more common in men.^41^ However, much less is known about sex differences in immune cells between CAD cases and controls. Here, we show a surprising male-to-female difference in upregulated genes in CD4 T cells. There is a group of genes that are equally important in males and females (ranked in the top 10): KLF2, CD3D and E, IL32, CD48 and CD52. The importance of CD3D and E suggests that the signaling strength of the TCR is increased in males and females with CAD. IL-32 is a known atherosclerosis genes.^27,42^ KLF2 is involved in Treg development,^43^ but the detailed function of KLF2 in CD4T cells in the development of CAD has not been reported. CD48 on T cells enhances TCR signaling through cis interactions with CD2, LAT and Lck.^44,45^, and is related to CD4T cell activation in other disease.^46,47^ Human and mouse antigen-activated T cells with high expression of CD52 suppressed other T cells.^48^ In DM-prone mice, transfer of lymphocyte populations depleted of CD52hi cells resulted in a substantially accelerated onset of DM.^48^ Many of the genes that are very important in predicting CAD in females are in the interferon pathway (IFITM2, IFITM3). Genes that are important in men but not women are TCF7, a transcription factor mainly expressed in naïve T cells, GNAI2 and lymhotoxin B (LTB).

The large difference in gene expression in CD4+ T cells between men and women was unexpected. Sex hormones are known to interact with the immune system on multiple levels. Autoimmune diseases are more common in women, a phenomenon also partly attributed to sex hormones.^49^ The expressions of several immune response genes in human PBMCs including *GATA3, IFNG, IL1B, LTA, NFKB1, PDCD1, STAT3, STAT5A, TBX21, TGFB1, TNFA* changes during menstrual cycle.^50^ In this present study, we showed that many immune response genes were highly expressed in women, along with many immune response signaling pathways. Furthermore, we showed from ML analysis that many immune response genes like IFITM2 and IFITM3, or IL32 are important to separate CAD vs non-CAD in female. Moreover, higher levels of IFNα2 were detected in female COVID-19 patients than in male patients in some cohort.^51^

Despite the decades-old knowledge that DM is a major risk factor for CVD, the reasons for this association are only partially understood. Interestingly, DM-accelerated atherosclerosis seems to be a human phenomenon and is not reproduced well in mouse models of atherosclerosis.^52^ We show a significant overlap in gene expression in subjects with CAD and DM, suggesting that the same genes may be important in both pathologies.

scRNA-Seq studies in PBMCs are attractive, because PBMCs are collected in many clinical studies, and the data analysis generates a wealth of information. scRNA-Seq has been applied to human PBMCs in atherosclerosis,^53,54^ and one other study combined single cell transcriptomes with surface marker expression (CITE-seq^55,56^). The present study is the largest scRNA-Seq and CITE-Seq study in CD4+ T cells. We found significant changes in cell proportions and gene expression patterns in subjects with DM or CAD. As discussed above, some of the key CAD driver genes seem are sex-specific, but most of the CAD driver genes are similar in subjects with and without DM.

TCF7 was strongly and commonly upregulated in CAD and DM, especially in men. The association of the TCF7 locus with T1DM is known^29^ but the function of TCF7 has not been reported either in CAD and or in DM. TCF7 encodes Transcription factor 7, also known as T-cell-specific transcription factor-1 (TCF-1). TCF7 along with LEF1 acts redundantly to control the maintenance and functional specification of Treg subsets to prevent autoimmunity.^57^ This current study suggested that this gene may connect CAD and DM.

The cell proportions of some clusters including cluster 8 (CCR2+Em), 12 (*MMP9*+), and 13 (PDL1+Em) were lower in females than males. Among them, CCR2+Em is a stable population of memory CD4+ T cells equipped for rapid recall response.^58^ The role of CCR2 in CD4T cells in context of CVD has not been elucidated,^12^ but decreased atherosclerotic lesion formation in *Ccr2*^*−/−*^ mice was reported.^59^ The lower proportion of these cells may be related to the (relative) atheroprotective role in women at least in older age (in this study, the age of women was 67 years ±8.04 years (mean±SD)).

We showed that CD4T cell interactions with myeloid, PSGL-1-L-selectin were stronger in males compared to females. L-selectin on T cells can interact with PSGL-1 on myeloid cells (where it is properly glycosylated, ref) and thus mediate secondary T cell tethering, which occurs when a freely flowing leukocyte transiently interacts with a rolling or adherent leukocyte or adherent leukocyte fragments and subsequently rolls on the endothelium.^60^ Indeed, differential expression or glycosylation of PSGL-1 in different leukocytes mediates selective recruitment of different subsets of monocytes or lymphocytes to atherosclerotic arteries.^61^ PSGL-1-expressing cytotoxic CD4 T cells are abundant in perimenopausal women with low estradiol levels contributes to T cell-mediated atherosclerotic development along with endothelial cell apoptosis,^62^ but this PSGL-1 is not glycosylated correctly to interact with L-selectin.^63^

A strength of the present study is that we included well matched males and females, DM+ and DM- and CAD+ and CAD-participants, obtained large number of cells, and analyzed surface phenotype plus transcriptome. A limitations of the present study is that it does not have a validation cohort (yet). The low number of participants not treated with statins means that most of the CAV participants were treated to the standard of care. Interestingly, all CAD+ subjects not on statins were males. It is known that statin compliance is lower in males than in females. This is a hypothesis-generating study; hypothesis testing remains to be done in mouse experiments.

## CONCLUSIONS

This study of CD4+ T cells provides strong evidence for overlapping gene expression patterns and pathways between CAD and DM, and strong sex differences.

## ACKNOWLDGEMENTS

R.S., J.V., and K.L. designed the study. C.A.M., and F.D. collected samples and data for the angiography analysis for calculating the score. R.S., J.V., and F.D. analyzed clinical data. R.S., J.V., J.M., and C.P.D. ran the scRNA-Seq experiments. R.S., J.V., A.F., P.R., W.P., T.P., C.A.M., A.S., L.L.L., C.C.H., and K.L. analyzed the data. R.G., S.S.A.S., V.S., A.A., and Y.G. conducted the bioinformatics analysis. R.S., and K.L. wrote the manuscript.

