## Supplemental_Materials for "Sex differences in coronary artery disease and diabetes revealed by scRNA-Seq and CITE-Seq of human CD4+ T cells"

Klaus Ley, MD

La Jolla Institute for Immunology

9420 Athena Circle

La Jolla, CA 92037, USA

(858) 752-6661 (tel)

(858) 752-6985 (fax)

Content

Supplemental Table 1-7 (Table S1-7)

Supplemental Figure 1-5 (Figure S1-5)

Legends of Supplemental Excel File 1-4

**Table S1. All reagents, manufacturers and catalogue numbers**

| **Reagent** | **Vendor** | **Catalogue #** |
| --- | --- | --- |
| cRPMI^*1^ |  | - |
| PBS | Fisher Scientific | 10010049 |
| FBS | Gemini | 100106-500mL |
| Trypan Blue | Thermo Fisher | 15250061 |
| Fc Block | BD | 564220 |
| SMK | BD | 633781 |
| DRAQ7 | BD | 564904 |
| Calcein AM^*2^ | Thermo Fisher | C1430 |
| DMSO | Thermo Fisher | D12345 |
| AbSeq | BD | - |
| Rhapsody Reagent Kit |  | 633771 |
| Rhapsodu Human Immune Response Panel |  | 633750 |
| Rhapsody Custom Reagent Panel |  | 633742 |
| AMPure XP beads | Beckman Coulter | A63881 |
| Ethanol | Milipore Sigma | E7023-500ml |
| Nuclease Free Water | Qiagen | 129114 |
| D1000 ScreenTape | Agilent | 5067-5584 |
| D1000 Sample Buffer | Agilent | 5067-5603 |
| Qubit Reagent Kit | Thermo Fisher | Q33231 |
| NovaSeq S1 100 Cycle Kit | Illumina | 20012865 |
| NovaSeq S2 100 Cycle Kit | Illumina | 20012862 |

*1: From 500mL of RPMI-1640 (with L-Glutamine) remove 75.5mL and transfer to two 50mL conical tubes for future potential use. Add 50 mL of 100% Human Serum Albumin (HSA). Add 5mL of the following (concentrations indicated are of stock solutions): HEPES (1M), Sodium pyruvate (100X) MEM-NEAA (100X), Pen Strep (Penicillin-Streptomycin), GlutaMAX Add 0.5mL of Mercaptoethanol (1000X). Mix by inversion in RPMI-1640 original bottle. Carefully transfer solution to a 500mL CorningStore at 4°C. Vacuum filter system with a 0.45mm filter size. *2: Resuspended in DMSO.

**Table S2. The viability of each sample tube**

| **Sample #** | **Average** | **Sample #** | **Average** |
| --- | --- | --- | --- |
| 1 | 92.50% | 34 | 77.30% |
| 2 | 90.00% | 36 | 91.38% |
| 3 | 84.00% | 37 | 90.79% |
| 4 | 89.00% | 38 | 87.86% |
| 6 | 93.00% | 39 | 84.09% |
| 7 | 90.50% | 40 | 90.06% |
| 8 | 93.00% | 41 | 90.09% |
| 9 | 90.00% | 42 | 95.53% |
| 10 | 94.79% | 43 | 93.23% |
| 11 | 91.75% | 44 | 87.69% |
| 12 | 93.10% | 45 | 97.40% |
| 13 | 94.47% | 46 | 95.17% |
| 14 | 89.55% | 47 | 84.85% |
| 15 | 84.97% | 48 | 88.61% |
| 16 | 92.13% | 49 | 92.62% |
| 17 | 93.62% | 50 | 92.39% |
| 18 | 90.33% | 51 | 89.20% |
| 19 | 90.72% | 52 | 94.69% |
| 20 | 93.14% | 53 | 89.34% |
| 21 | 91.92% | 54 | 93.21% |
| 22 | 93.43% | 55 | 90.28% |
| 23 | 90.45% | 56 | 93.56% |
| 24 | 94.54% | 57 | 98.80% |
| 25 | 90.02% | 59 | 95.65% |
| 26 | 88.50% | 60 | 94.42% |
| 27 | 82.70% | 61 | 93.10% |
| 28 | 92.47% | 62 | 93.61% |
| 29 | 89.52% | 64 | 95.96% |
| 30 | 88.70% | 65 | 83.35% |
| 31 | 89.74% | **Min** | 77.1% |
| 32 | 77.10% | **Max** | 98.8% |
| 33 | 89.26% | **Median** | 90.8% |
|  |  | **Average** | 90.7% |

**Table S3. The information of 51 AbSeq antibodies**

| **Specificity** | **Clone** | **Catalogue Number** |
| --- | --- | --- |
| CD11b | M1/70 | 940008 |
| CD11c | B-LY6 | 940024 |
| CD123 (IL-3RA) | 7G3 | 940020 |
| CD126 (IL-6R) | M5 | 940090 |
| CD127 (IL-7R) | HIL-7R-M21 | 940012 |
| CD137 | 4B4-1 | 940055 |
| CD14 | MPHIP9 | 940005 |
| CD141 | 1A4 | 940079 |
| CD142 | HTF-1 | 940280 |
| CD152 (CTLA-4) | BNI3 | 940034 |
| CD154 | TRAP1 | 940053 |
| CD16 | 3G8 | 940006 |
| CD163 | GHI/61 | 940058 |
| CD183 (CXCR3) | 1C6/CXCR3 | 940030 |
| CD184 (CXCR4) | 12G5 | 940056 |
| CD185 (CXCR5) | RF8B2 | 940042 |
| CD19 | SJ25C1 | 940004 |
| CD192 (CCR2) | 1D9 | 940286 |
| CD194 (CCR4) | 1G1 | 940047 |
| CD195 (CCR5) | 2D7/CCR5 | 940050 |
| CD196 (CCR6) | 11A9 | 940033 |
| CD197 (CCR7) | 3D12 | 940014 |
| CD2 | RPA-2.10 | 940046 |
| CD20 | 2H7 | 940016 |
| CD206 | 19.2 | 940068 |
| CD223 (LAG-3) | T47-530 | 940080 |
| CD25 | 2A3 | 940009 |
| CD27 | M-T271 | 940018 |
| CD3 | SK7 | 940000 |
| CD36 | CB38 (NL07) | 940224 |
| CD38 | HIT2 | 940013 |
| CD4 | SK3 | 940001 |
| CD45RA | HI100 | 940011 |
| CD45RO | UCHL1 | 940022 |
| CD56 | NCAM16.2 | 940007 |
| CD69 | FN50 | 940019 |
| CD8 | RPA-T8 | 940003 |
| CD86 | 2331(FUN-1) | 940025 |
| CD9 | M-L13 | 940078 |
| HLA-DR (CD74) | G46-6 | 940010 |
| CD24 | ML5 | 940028 |
| CD33 | WM53 | 940031 |
| IgM | G20-127 | 940276 |
| IgD | IA6-2 | 940026 |
| CD43 | 1G10 | 940728 |
| CD273 | MIH18 | 746072 |
| CD274 | MIH1 | 940035 |
| CD95 | DX2 | 940037 |
| CD279 | MIH4 | 940467 |
| TLR4-APC | 610015 | not applicable |
| SLAN-PE | M-DC8 | not applicable |
| PE | E31-1459 | 460077 |
| APC | E30-221 | 460078 |

**Table S4. Thresholds of each antibody expression.**

| **Antibody** | **threshold** | **Antibody** | **threshold** |
| --- | --- | --- | --- |
| **CD2** | 2.4 | **CD126** | not use for clustering |
| **CD3** | 2 | **CD127** | 0.8 |
| **CD4** | 1.65 | **CD137** | not use for clustering |
| **CD8** | 3.7 | **CD141** | 0.5 |
| **CD9** | 1.3 | **CD142** | not use for clustering |
| **CD11b** | remove from all the analysis | **CD152 CTLA4** | not use for clustering |
| **CD11c** | 2 | **CD154** | remove from all the analysis |
| **CD14** | 1.65 | **CD163** | not use for clustering |
| **CD16** | 3.25 | **CD183** | 1.1 |
| **CD19** | 1.3 | **CD184** | not use for clustering |
| **CD20** | 0.45 | **CD185** | 0.2 |
| **CD24** | 0.6 | **CD192** | 1.25 |
| **CD25** | not use for clustering | **CD194** | not use for clustering |
| **CD27** | 1.15 | **CD195** | not use for clustering |
| **CD33** | 1.3 | **CD196** | not use for clustering |
| **CD36** | 3.4 | **CD197** | not use for clustering |
| **CD38** | 1 | **CD206** | not use for clustering |
| **CD43** | 2 | **CD223** | not use for clustering |
| **CD45RA** | 2.15 | **CD273 PDL2** | not use for clustering |
| **CD45RO** | 0.8 | **CD274 PDL1** | 1 |
| **CD56** | 0.5 | **CD279 PD1** | 0.75 |
| **CD69** | not use for clustering | **CD74** | 1.6 |
| **CD86** | 0.9 | **IgD** | 1.5 |
| **CD95** | 1.85 | **IgM** | 0.8 |
| **CD123** | 1 | **SA06 TLR4** | 0.1 |
|  |  | **SA16 SLAN** | 0.37 |

**Table S5A-E. Clinical statistics of all the samples.**

**A. CAD Low vs CAD High in Diabetic/Non-Diabetic Men on Statins and Not Diabetic Men not on Statins.**

|  | **Diabetic Men on Statins (n=9/8 per group)** | | | **Non-Diabetic Men on Statins (n=9/8 per group)** | | | **Not Diabetic Men not on Statins** | | |
| --- | --- | --- | --- | --- | --- | --- | --- | --- | --- |
|  |  |  |  |  |  |  | **(n=5 per group)** | | |
| **Variable** | **CAD Low** | **CAD High** | **P-value**^†^ | **CAD Low** | **CAD High** | **P-value**^†^ | **CAD Low** | **CAD High** | **P-value**^†^ |
| Count [%] or Median [± SD] | **Gensini  < 6** | **Gensini > 30** |  | **Gensini  < 6** | **Gensini  > 30** |  | **Gensini < 6** | **Gensini > 30** |  |
|  | (n=9) | (n=8) |  | (n=8) | (n=9) |  | (n=5) | (n=5) |  |
| **General Characteristics** | |  |  |  |  |  |  |  |  |
| Age (years) | 62 [±8.48] | 63 [±10.29] | 0.7 | 65 [±6.23] | 64 [±6.73] | 0.92 | 63 [±12.97] | 64 [±13.40] | 0.92 |
| Race (Caucasian) | 9 [100%] | 8 [100%] |  | 8 [100%] | 8 [88.9%] | 0.33 | 3 [60%] | 5 [100%] | 0.12 |
| Ethnicity (Non-Hispanic) | 9 [100%] | 8 [100%] |  | 8 [100%] | 7 [77.78%] | 0.15 | 4 [80%] | 5 [100%] | 0.29 |
| Current Smoker (Yes) | 0 [0%] | 0 [0%] |  | 0 [0%] | 0 [0%] |  | 0 [0%] | 0 [0%] |  |
| Former Smoker (Yes) | 5 [55.6%] | 5 [62.5%] | 0.77 | 4 [50%] | 3 [33.3%] | 0.49 | 3 [60%] | 3 [60%] |  |
| BMI | 35 [±6.51] | 32 [±6.86] | 0.54 | 32 [±7.61] | 31 [±5.76] | 0.51 | 30 [±5.13] | 29 [±5.72] | 0.68 |
| BP Systolic | 147 [±19.76] | 132 [±18.99] | 0.12 | 138 [±18.96] | 141 [±5.94] | 0.33 | 123 [±13.83] | 143 [±12.63] | 0.07 |
| BP Diastolic | 80 [±10.49] | 70 [±12.67] | 0.12 | 81 [±15.54] | 78 [±13.13] | 0.6 | 79 [±13.12] | 83 [±17.08] | 0.92 |
| **Medications** |  |  |  |  |  |  |  |  |  |
| Diuretics (Yes) | 0 [0%] | 2 [25%] | 0.11 | 1 [12.5%] | 4 [44.4%] | 0.15 | 0 [0%] | 1 [20%] | 0.29 |
| Beta Blockers (Yes) | 7 [77.8%] | 5 [62.5%] | 0.49 | 4  [50%] | 4 [44.4%] | 0.82 | 2 [40%] | 2 [40%] |  |
| Calcium Channel Blockers (Yes) | 1 [11.1%] | 1 [12.5%] | 0.93 | 2 [25%] | 2 [22.2%] | 0.89 | 0 [0%] | 0 [0%] |  |
| ACE (Yes) | 4 [44.4%] | 4 [50%] | 0.82 | 2 [25%] | 2 [22.2%] | 0.89 | 1 [20%] | 1 [20%] |  |
| ATR (Yes) | 2 [22.2%] | 1 [12.5%] | 0.6 | 1 [12.5%] | 0 [0%] | 0.27 | 1 [20%] | 0 [100%] | 0.29 |
| NSAID (Yes) | 8 [88.9%] | 7 [87.5%] | 0.93 | 7 [87.5%] | 9 [100%] | 0.27 | 3 [60%] | 4 [80%] | 0.49 |
| **Lab Values** |  |  |  |  |  |  |  |  |  |
| Creatinine | 0.66 [±0.35] | 0.85 [±0.17] | 0.34 | 0.95 [±0.18] | 0.99 [±0.24] | 0.85 | 0.84 [±0.15] | 0.82 [±0.11] | 0.99 |
| Hs-CRP | 1.54 [±1.30] | 2.18 [±1.98] | 0.48 | 2.25 [±2.45] | 2.98 [±3.34] | 0.82 | 4.06 [±5.60] | 1.77 [±2.14] | 0.84 |
| Total Cholesterol (mg/dL) | 134 [±30.62] | 135 [±35.38] | 0.96 | 129 [±22.12] | 140 [±40.03] | 0.6 | 150 [±27.33] | 169 [±57.09] | 0.84 |
| Triglyceride (mg/dL) | 146 [±100.08] | 162 [±94.74] | 0.54 | 125 [±53.86] | 103 [±64.02] | 0.43 | 84 [±39.23] | 89 [±12.46] | 0.43 |
| HDL Cholesterol (mg/dL) | 38 [±9.42] | 36 [±10.31] | 0.6 | 39 [±15.82] | 43 [±10.39] | 0.33 | 42 [±8.53] | 40 [±8.35] | 0.76 |
| LDL Cholesterol (mg/dL) | 72 [±28.29] | 72 [±31.61] | 0.89 | 69 [±10.97] | 80 [±29.89] | 0.6 | 94 [±24.97] | 114 [±48.43] | 0.69 |
| Glucose (mg/dL) | 151 [±60.02] | 144 [±36.64] | 0.74 | 98 [±11.93] | 94 [±13.36] | 0.81 | 95 [±13.87] | 101 [±6.91] | 0.76 |
| A1c (%) | 7.31 [±1.46] | 7.18 [±1.31] | 0.81 | 5.81 [±0.76] | 5.54 [±0.44] | 0.85 | 5.50 [±0.48] | 5.67 [±0.17] | 0.92 |
| **Disease Severity** | |  |  |  |  |  |  |  |  |
| Gensini Scores | 1.83 [±1.97] | 66.19 [±31.75] | 0 | 3.00 [±2.67] | 70.33 [37.52] | 0 | 1.10 [±2.46] | 81.80 [±32.45] | 0.03 |

†Categorical variables were calculated by Chi-square test and continuous variables by Mann Whitney

**B. CAD Low vs CAD High in Women on Statins with and without diabetes.**

| **Variable** | **CAD Low** | **CAD High** | **P-value**^†^ |
| --- | --- | --- | --- |
| Count [%] or Median [± SD] | **Gensini < 6** | **Gensini > 30** |  |
|  | (n=7) | (n=9) |  |
| **General Characteristics** | | | |
| Age (years) | 65 [±7.72] | 67 [±8.82] | 0.47 |
| Race (Caucasian) | 7 [100%] | 9 [100%] | 0.24 |
| Ethnicity (Non-Hispanic) | 6 [85.7%] | 9 [100%] | 0.24 |
| Diabetes (Yes) | 4 [57.1%] | 5 [55.6%] | 0.95 |
| Current Smoker (Yes) | 2 [28.6%] | 3 [33.3%] | 0.84 |
| Former Smoker (Yes) | 4 [57.1%] | 2 [22.2%] | 0.15 |
| BMI | 39 [±4.94] | 29 [±6.26] | 0.03 |
| BP Systolic | 124 [±16.78] | 150 [±26.29] | 0.08 |
| BP Diastolic | 71 [±10.92] | 77 [±14.67] | 0.44 |
| **Medications** |  |  |  |
| Diuretics (Yes) | 3 [42.86%] | 4 [44.4%] | 0.94 |
| Beta Blockers (Yes) | 5 [71.43%] | 5 [55.6%] | 0.52 |
| Calcium Channel Blockers (Yes) | 1 [14.3%] | 5 [55.6%] | 0.09 |
| ACE (Yes) | 4 [57.14%] | 4 [44.4%] | 0.61 |
| ATR (Yes) | 0 [100%] | 0 [100%] |  |
| NSAID (Yes) | 5 [71.4%] | 7 [77.8%] | 0.77 |
| **Lab Values** |  |  |  |
| Creatinine | 0.81 [±0.13] | 0.70 [±0.17] | 0.18 |
| Hs-CRP | 6.25 [±4.14] | 3.17 [±2.42] | 0.07 |
| Total Cholesterol (mg/dL) | 159 [±19.70] | 163 [±55.38] | 0.53 |
| Triglyceride (mg/dL) | 178 [±69.92] | 101 [±49.50] | 0.05 |
| HDL Cholesterol (mg/dL) | 41 [±10.09] | 54 [±20.35] | 0.17 |
| LDL Cholesterol (mg/dL) | 88 [±21.58] | 92 [±41.28] | 0.99 |
| Glucose (mg/dL) | 128 [±40.23] | 111 [±23.31] | 0.53 |
| A1c (%) | 7.3 [±2.12] | 6.36 [±0.62] | 0.68 |
| **Disease Severity** | | | |
| Gensini Scores | 3.00 [±2.81] | 39.22 [±7.55] | 0.005 |

**C. CAD Low vs CAD High in all the participants**

| Variable Count [%] or Mean [± SD] | **CAD Low (n=29)** | **CAD High (n=32)** | **p-value** |
| --- | --- | --- | --- |
| **Demographics** | | | |
| Age (years) | 64 [±8.27] | 65 [±9.12] | 0.41 |
| Sex (Male) | 22 (76%) | 22 (69%) | 0.54 |
| Race (Caucasian) | 27 (93%) | 31 (97%) | 0.50 |
| Ethnicity (Non-Hispanic) | 27 (93%) | 30 (94%) | 0.92 |
| Diabetes (Yes) | 13 (45%) | 14 (44%) | 0.93 |
| Smoking | 18 (62%) | 16 (50%) | 0.34 |
| BMI | 34.2 [±6.65] | 30.6 [±6.02] | 0.04 |
| BP Systolic | 135 [±19.85] | 143 [±19.36] | 0.09 |
| BP Diastolic | 78 [±12.56] | 76 [±13.97] | 0.57 |
| **Medications** | | | |
| Statins (Yes) | 24 (83%) | 26 (81%) | 0.88 |
| Diuretics (Yes) | 4 (14%) | 11 (34%) | 0.06 |
| Beta Blockers (Yes) | 18 (62%) | 16 (50%) | 0.34 |
| Calcium Channel Blockers (Yes) | 4 (14%) | 9 (28%) | 0.17 |
| ACE (Yes) | 11 (38%) | 11 (34%) | 0.77 |
| ATR (Yes) | 4 (14%) | 2 (6%) | 0.32 |
| NSAID (Yes) | 23 (79%) | 28 (88%) | 0.39 |
| **Lab Values** | | | |
| Creatinine | 0.81 [±0.25] | 0.84 [±0.20] | 0.84 |
| Hs-CRP | 3.3 [±3.71] | 2.64 [±2.48] | 0.78 |
| Total Cholesterol (mg/dL) | 141 [±27.01] | 152 [±46.99] | 0.65 |
| Triglyceride (mg/dL) | 137 [±76.39] | 114 [±67.66] | 0.19 |
| HDL Cholesterol (mg/dL) | 40 [±11.05] | 44 [±14.73] | 0.36 |
| LDL Cholesterol (mg/dL) | 79 [±23.49] | 89 [±38.98] | 0.60 |
| Glucose (mg/dL) | 121 [±44.74] | 113 [±29.46] | 0.98 |
| A1c (%) | 6.6 [±1.56] | 6.2 [±0.98] | 0.81 |
| **Disease Severity** | | | |
| Gensini Scores | 2.3 [±2.46] | 61.3 [±31.66] | <.0001 |

**D. Diabetes vs No Diabetes in all the participants**

| Variable Count [%] or Mean [± SD] | Diabetes (n=27) | No Diabetes (n=34) | p-value |
| --- | --- | --- | --- |
| **Demographics** | | | |
| Age (years) | 65 [±8.46] | 64 [±8.98] | 0.87 |
| Sex (Male) | 17 (63%) | 27 (79%) | 0.15 |
| Race (Caucasian) | 27 (100%) | 31 (91%) | 0.11 |
| Ethnicity (Non-Hispanic) | 27 (100%) | 30 (88%) | 0.07 |
| Smoking (Yes) | 17 (63%) | 17 (50%) | 0.59 |
| BMI | 33.5 [±6.46] | 31.4 [±6.53] | 0.25 |
| BP Systolic | 142 [±24.62] | 136 [±14.91] | 0.52 |
| BP Diastolic | 75 [±11.66] | 79 [±14.23] | 0.31 |
| **Medications** | | | |
| Statins (Yes) | 26 (96%) | 24 (71%) | 0.01 |
| Diuretics (Yes) | 6 (22%) | 9 (26%) | 0.70 |
| Beta Blockers (Yes) | 18 (67%) | 16 (47%) | 0.13 |
| Calcium Channel Blockers (Yes) | 6 (22%) | 7 (21%) | 0.88 |
| ACE (Yes) | 13 (48%) | 9 (26%) | 0.08 |
| ATR (Yes) | 4 (15%) | 2 (6%) | 0.24 |
| NSAID (Yes) | 24 (89%) | 27 (79%) | 0.32 |
| **Lab Values** | | | |
| Creatinine | 0.73 [±0.24] | 0.90 [±0.18] | 0.007 |
| Hs-CRP | 8.3 [±28.42] | 3.1 [±3.59] | 0.48 |
| Total Cholesterol (mg/dL) | 143 [±36.43] | 150 [±40.95] | 0.52 |
| Triglyceride (mg/dL) | 153 [±87.38] | 103 [±48.21] | 0.03 |
| HDL Cholesterol (mg/dL) | 40 [±14.77] | 43 [±11.80] | 0.16 |
| LDL Cholesterol (mg/dL) | 78 [±30.83] | 90 [±33.60] | 0.26 |
| Glucose (mg/dL) | 141 [±43.71] | 98 [±14.15] | <.0001 |
| A1c (%) | 7.3 [±1.41] | 5.7 [±0.50] | <.0001 |
| **Disease Severity** | | | |
| Gensini Scores | 29.4 [±32.84] | 36.28 [±40.95] | 0.69 |

**E. Men vs Women in all the participants**

| Variable Count [%] or Mean [± SD] | Men (n=44) | Women (n=17) | p-value |
| --- | --- | --- | --- |
| **Demographics** | | | |
| Age (years) | 64 [±8.86] | 67 [±8.04] | 0.24 |
| Race (Caucasian) | 41 (93%) | 17 (100%) | 0.27 |
| Ethnicity (Non-Hispanic) | 41 (93%) | 16 (94%) | 0.89 |
| Diabetes (Yes) | 17 (39%) | 10 (59%) | 0.15 |
| Smoking (Yes) | 23 (52%) | 11 (65%) | 0.38 |
| BMI | 32.9 [±6.33] | 33.4 [±7.10] | 0.37 |
| BP Systolic | 138 [±16.86] | 141 [±26.53] | 0.91 |
| BP Diastolic | 78 [±13.44] | 75 [±12.72] | 0.30 |
| **Medications** | | | |
| Statins (Yes) | 34 (77%) | 16 (94%) | 0.13 |
| Diuretics (Yes) | 8 (18%) | 7 (41%) | 0.06 |
| Beta Blockers (Yes) | 24 (55%) | 10 (59%) | 0.76 |
| Calcium Channel Blockers (Yes) | 6 (14%) | 7 (41%) | 0.02 |
| ACE (Yes) | 14 (32%) | 8 (47%) | 0.27 |
| ATR (Yes) | 5 (11%) | 1 (6%) | 0.52 |
| NSAID (Yes) | 38 (86%) | 13 (76%) | 0.35 |
| **Lab Values** | | | |
| Creatinine | 0.85 [±0.24] | 0.75 [±0.15] | 0.04 |
| Hs-CRP | 2.4 [±2.82] | 4.4 [±3.44] | 0.001 |
| Total Cholesterol (mg/dL) | 140 [±35.70] | 164 [±42.58] | 0.05 |
| Triglyceride (mg/dL) | 123 [±74.17] | 131 [±68.80] | 0.54 |
| HDL Cholesterol (mg/dL) | 39 [±10.68] | 48 [±16.93] | 0.06 |
| LDL Cholesterol (mg/dL) | 81 [±31.30] | 94 [±35.20] | 0.16 |
| Glucose (mg/dL) | 116 [±39.98] | 118 [±30.93] | 0.41 |
| A1c (%) | 6.3 [±1.21] | 6.8 [±1.45] | 0.06 |
| **Disease Severity** | | | |
| Gensini Scores | 36.8 [±42.13] | 24.1 [±19.05] | 0.58 |

**Table S6. The 31 overlapped genes significantly upregulated in both CAD and DM**

| LAIR2 | Leukocyte-Associated Immunoglobulin-Like Receptor 2, CD306 |
| --- | --- |
| LTB | lymphotoxin β |
| TCF7 | TCF1,exhaustion marker. necessary for T cell differentiation |
| GNAI2 | G Protein Subunit Alpha I2 |
| R3HDM4 | nucleic acid binding protein |
| CTSW | Cathepsin W, regulation of T-cell cytolytic activity |
| CD7 | immunoglobulin family, T-B cell interactions |
| STAT6 | Signal Transducer And Activator Of Transcription 6, IL-4 signaling |
| IFITM3 | IFN-induced antiviral protein, disrupts intracellular cholesterol homeostasis |
| TNFRSF25 | Death Receptor 3 (DR3) for TWEAK. Activates NF-kappa-B, apoptosis |
| PXN | paxillin, actin-membrane attachment at sites of cell adhesion |
| XPO6 | exportin6, mediates nuclear export of actin and profilin-actin complexes |
| SRGN | serglycin, cell granule proteoglycan |
| LGALS9 | Galectin 9, S-type lectin for β-Gal, binds HAVCR2 to induce Th1 death |
| S100A12 | binds calcium, zinc, copper. Binds to AGE receptor, pro-inflammatory |
| CD3E | εsubunit of CD3 |
| ZAP70 | TCR signaling, homologue of SYK |
| CD4 | co-receptor for MHC-II |
| TSPAN32 | tetraspanin associated with malignancies |
| ITGB2 | β2 integrin, CD18 |
| IGBP1 | immunoglobulin binding protein-1. regulates phosphatases PP2A, PP4, 6 |
| JUNB | JUN-B proto-oncogene, subunit of AP-1 |
| CD37 | tetraspanin, complexes with integrins, T-B cell interactions |
| CTSD | cathepsin D protease, activates hormones and growth factors |
| CD6 | adhesion molecule, binds ALCAM/CD166 |
| CD3G | γsubunit of CD3 |
| BIN2 | Modulates membrane curvature and mediates membrane tubulation |
| CD52 | CAMPATH-1 antigen, carries and orients glycans |
| CD74 | Stabilizes empty MHC class II alpha/beta heterodimers |
| CD27 | TNFRSF7. Required for generation and maintenance of T cell immunity |
| NKG7 | Natural Killer Cell Granule Protein 7, in granule membrane |

**Supplemental Figure 1.** Ridgeline plots of the unthresholded expressions of the 34 antibodies used for clustering for each main cell types [CD4 T cells, CD8 T cells, myeoid cells, NK cells and B cells using Seurat. These plots separately show the distribution of each value of CLR normalized antibody derived tag for each main cell type. Expression levels are shown on x-axis. Thresholding lines are shown in black line in each plot.

**Supplemental Figure 2.** Violin plots for surface marker expressions of each subcluster.

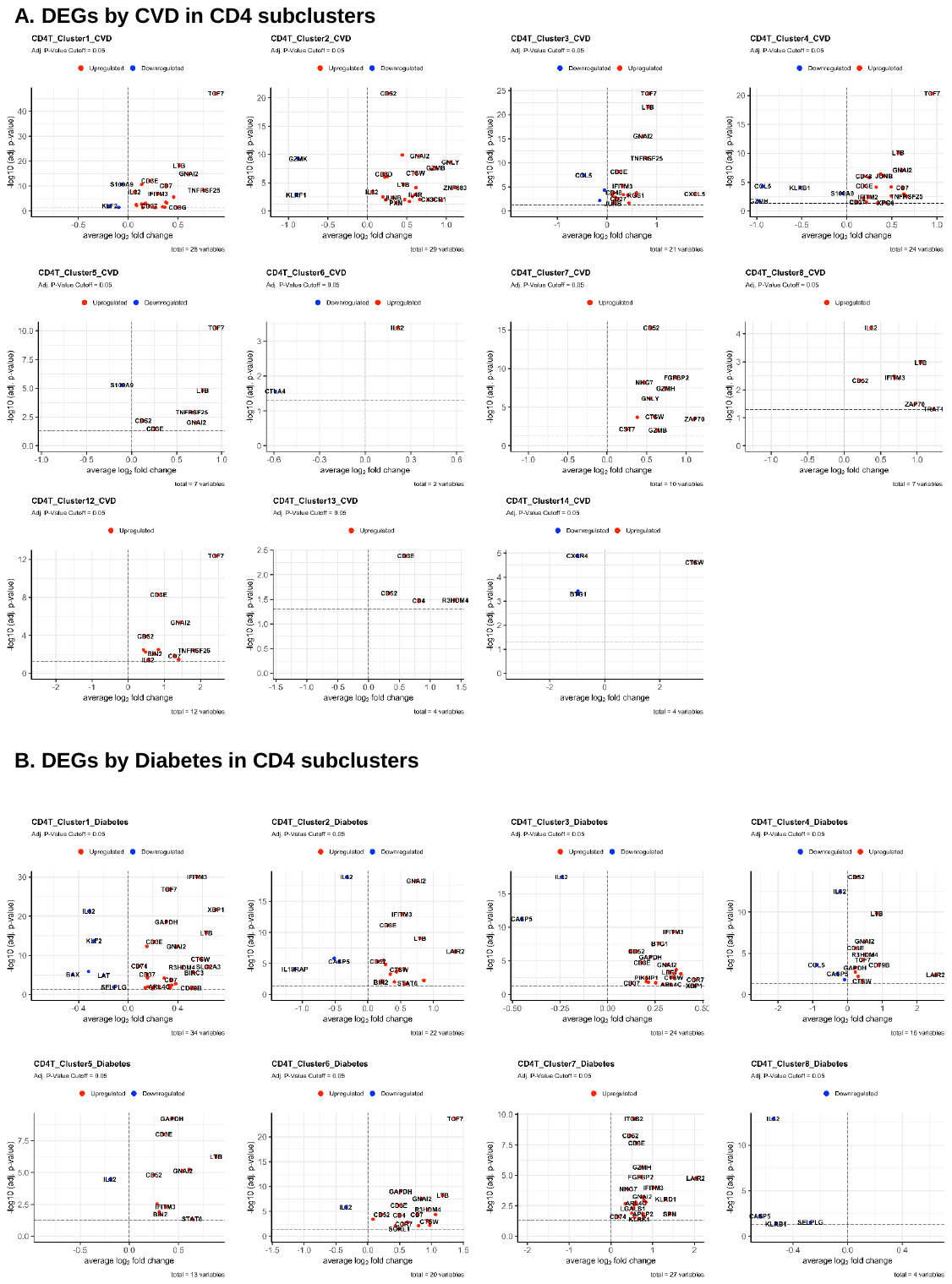

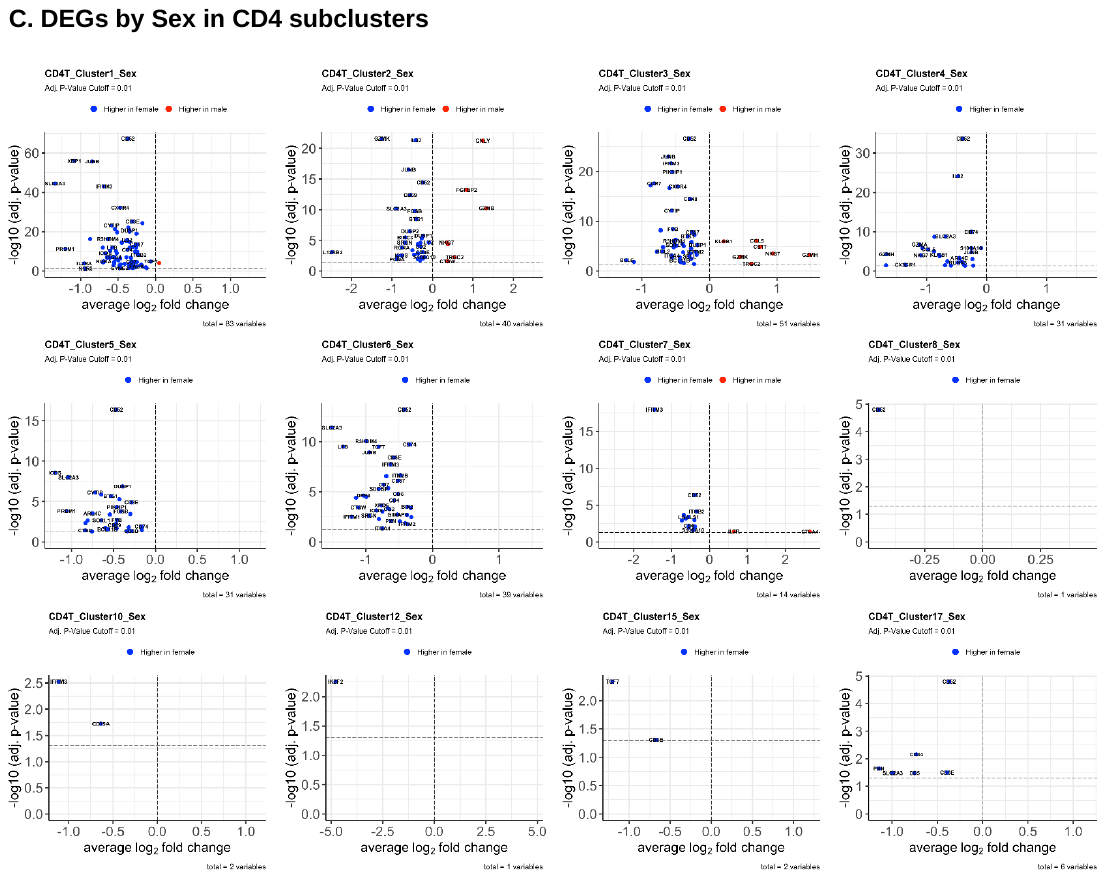

**Supplemental Figure 3.** Dotplots of differentially expressed genes between CAD+ vs CAD-, and between diabetes+ vs diabetes- in each CD4 T cell subcluster. The size of dots represents log(pct.1/pct.2). pct.1, the proportion of cells expressing each gene in Diabetes or CAD+; pct.2, the proportion of cells expressing each gene in Diabetes- or CAD-.

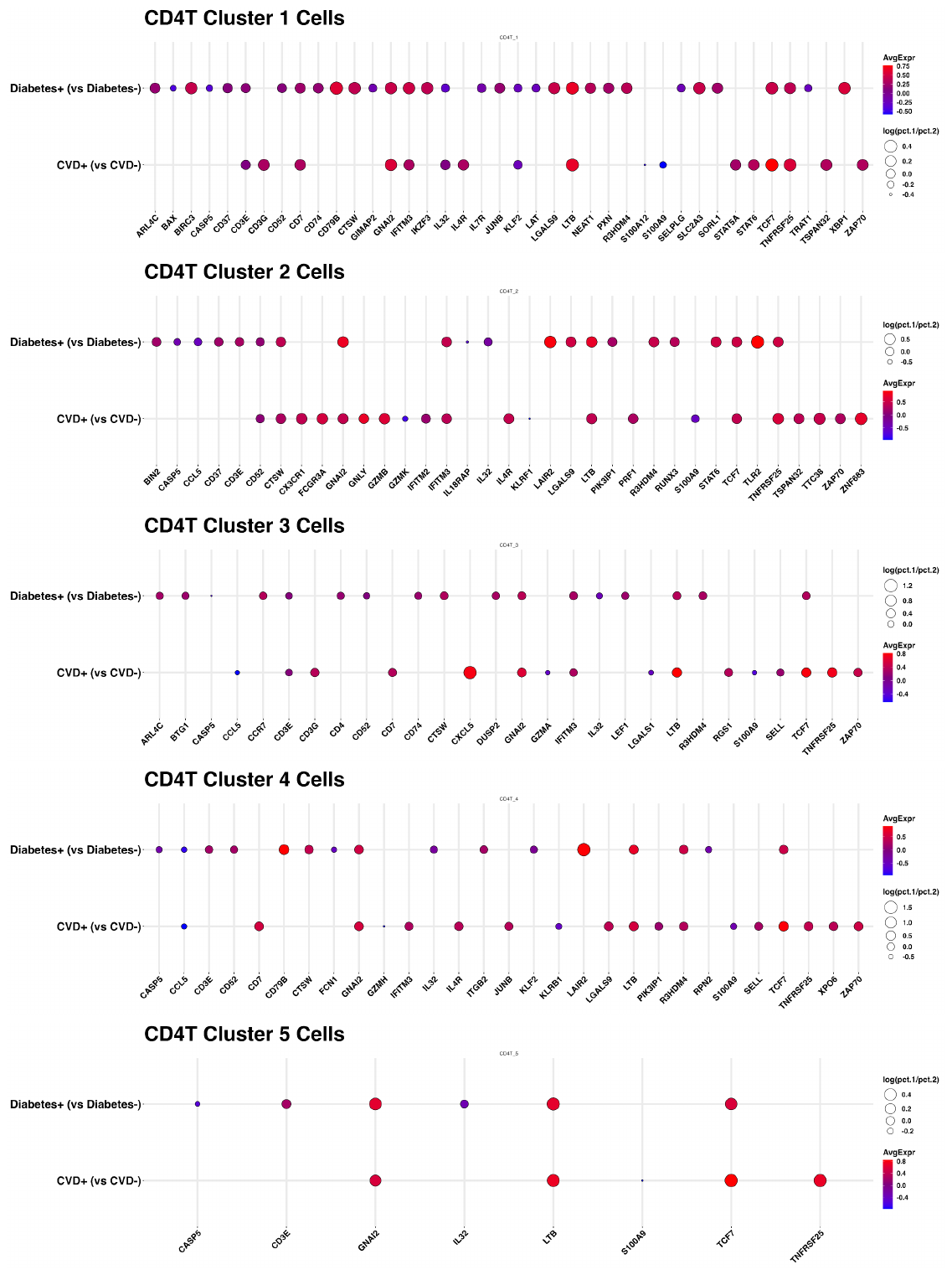

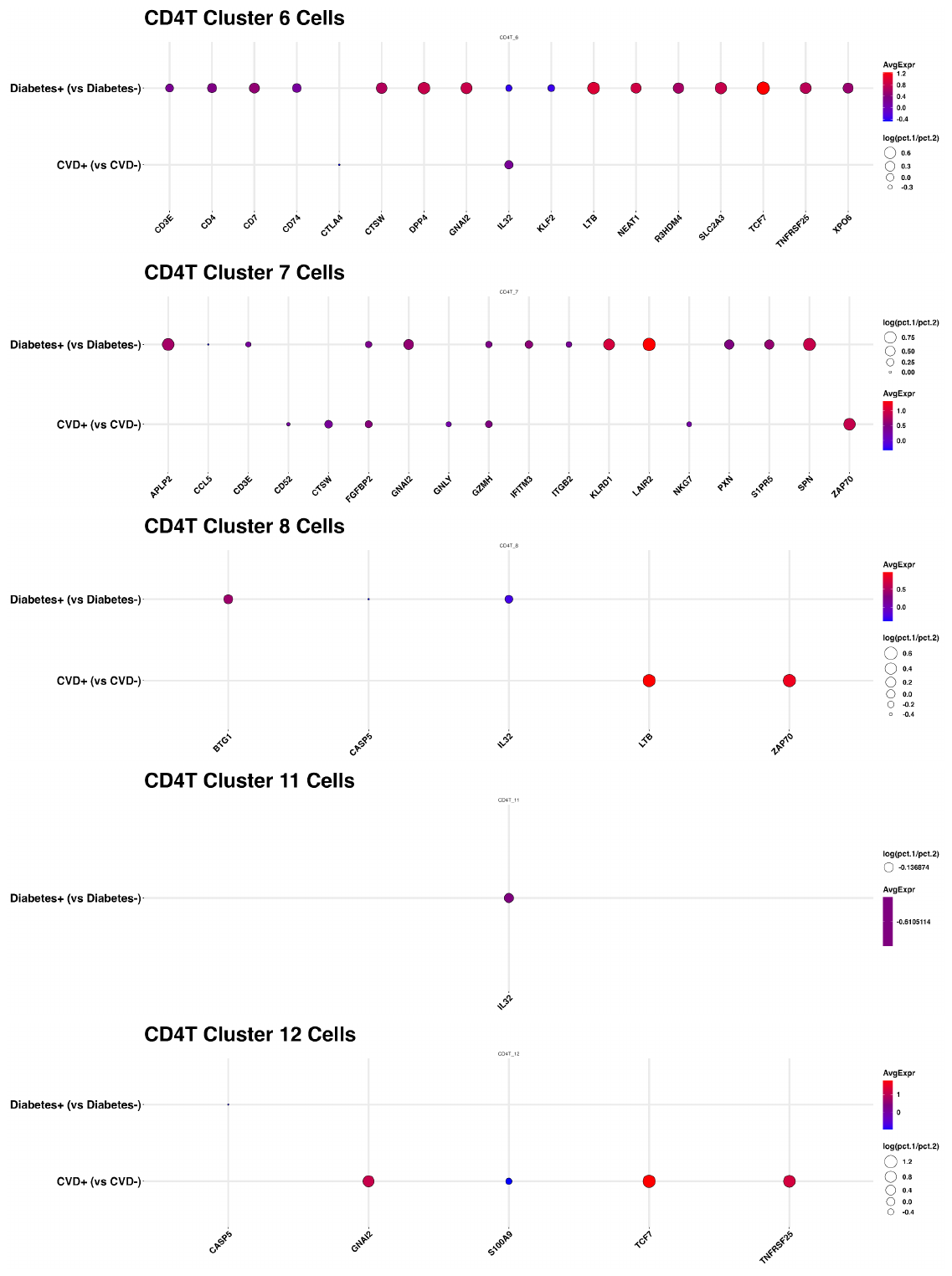

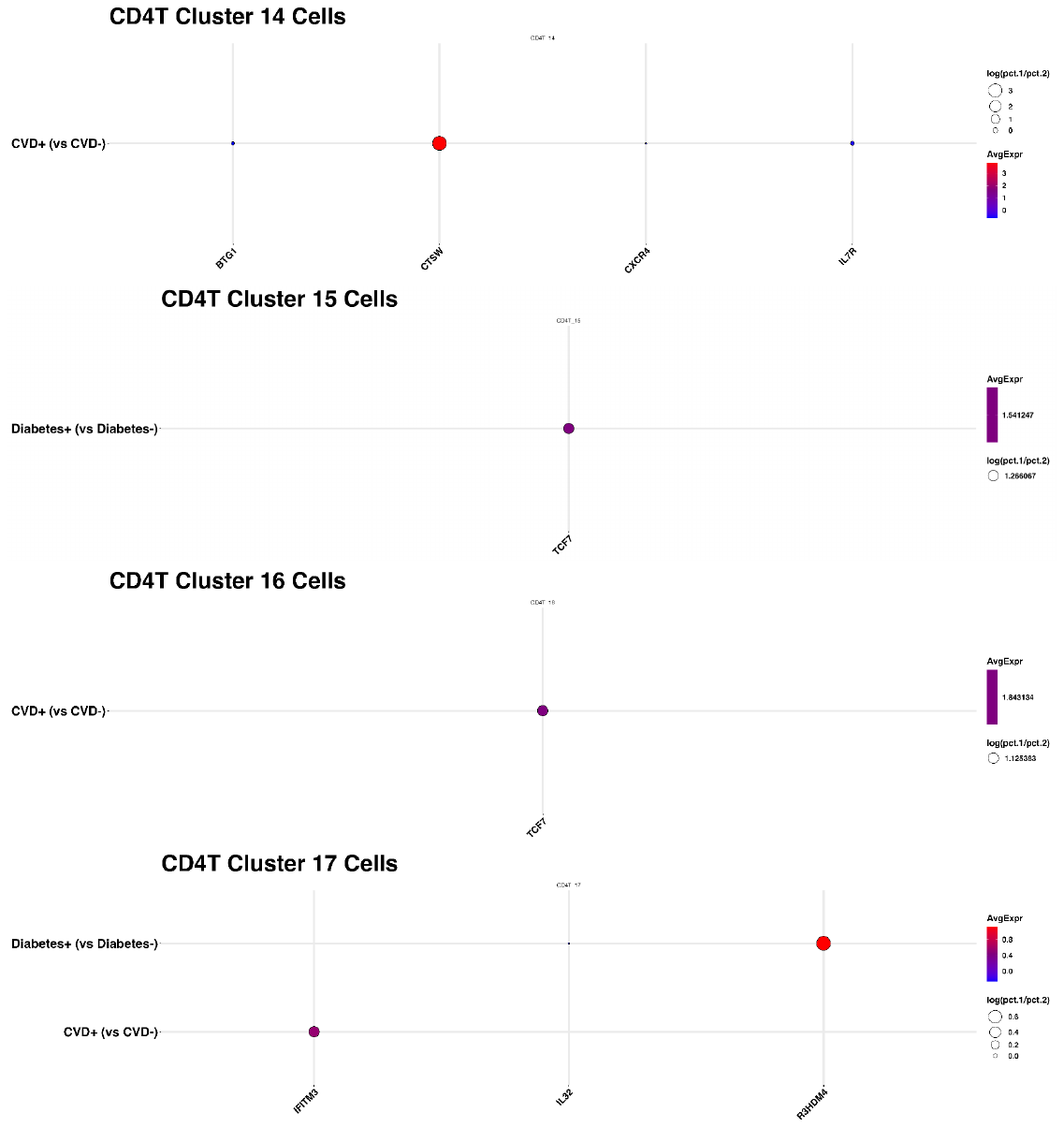

**Supplemental Figure 4.** A. Volcano plots for significantly upregulated genes by CAD in CD4 subclusters. B. Volcano plots for significantly upregulated genes by Diabetes in CD4 subclusters. Volcano plots for significantly upregulated genes by Sex in CD4 subclusters.

**Supplemental Figure 5.** Bubble plots of all interaction analysis between CD4T-CD4T cells and CD4T-myeloid cells with significant communication probabilities.

**Supplemental Excel File 1. All the counts of gene expression in each single cell.**

**Supplemental Excel File 2. Differentially expressed genes in each cluster against all the others.**

**Supplemental Excel File 3. Differentially expressed genes by disease status in total CD4 T cells.**

**Supplemental Excel File 4. Differentially expressed genes by disease status in each CD4T subcluster.**
